# Community assembly modeling of microbial evolution within Barrett’s esophagus and esophageal adenocarcinoma

**DOI:** 10.1101/2025.01.14.633020

**Authors:** Caitlin Guccione, Igor Sfiligoi, Antonio Gonzalez, Justin P. Shaffer, Mariya Kazachkova, Yuhan Weng, Daniel McDonald, Shailja C. Shah, Samuel S. Minot, Thomas Paulson, William M. Grady, Ludmil B. Alexandrov, Rob Knight, Kit Curtius

## Abstract

Mathematical modeling of somatic evolution, a process impacting both host cells and microbial communities in the human body, can capture important dynamics driving carcinogenesis. Here we considered models for esophageal adenocarcinoma (EAC), a cancer that has dramatically increased in incidence over the past few decades in Western populations, with high case fatality rates due to late-stage diagnoses. Despite advancements in genomic analyses of the precursor Barrett’s esophagus (BE), prevention of late-stage EAC remains a significant clinical challenge. Previous microbiome studies in BE and EAC have focused on quantifying static microbial abundance differences rather than evolutionary dynamics. Using whole genome sequencing data from esophageal tissues, we first applied a robust bioinformatics pipeline to extract non-host DNA reads, mapped these putative reads to microbial taxa, and retained those taxa with high genomic coverage. When applying mathematical models of microbial evolution to sequential stages of progression to EAC, we observed evidence of neutral dynamics in community assembly within normal esophageal tissue and BE, but not EAC. In a case-control study of BE patients who progressed to EAC cancer outcomes (CO) versus those who had non-cancer outcomes (NCO) during follow-up (mean=10.5 years), we found that *Helicobacter pylori* deviated significantly from the neutral expectation in BE NCO, suggesting that factors related to *H. pylori* or *H. pylori* infection itself may influence EAC risk. Additionally, simulations incorporating selection recapitulated non-neutral behaviors observed in the datasets. Formally modeling dynamics during progression holds promise in clinical applications by offering a deeper understanding of microbial involvement in cancer development.

## Introduction

To improve our understanding of microbiome dynamics in certain environments, we must elucidate the mechanisms driving microbial community assembly. The majority of studies toward this aim focus on the relative contribution of niche-vs. neutral processes. Mathematical modeling can be used to quantify such dynamics in microbial populations but thus far has been underutilized in the microbiome analysis of human cancers. The assembly dynamics of microbes in relation to human cancer progression, as well as their potential to enhance predictive models of cancer progression and metastasis, remain largely unexplored. The involvement of microbes in human cancer development is well-supported, specifically that many cancers (such as breast, lung, and colon cancers) evolve in the presence of microbial communities [1–4]. While research on the cancer microbiome is increasing, many sequencing studies of large patient cohorts to date have primarily focused on describing microbial abundances at static time points rather than modeling the temporal dynamics of populations or communities explicitly. Development of these models is essential for hypothesis testing, and for supporting rigorous conclusions on expected versus observed community assembly characteristics over time in the human microbiome.

Esophageal adenocarcinoma (EAC) is a cancer that occurs in the distal esophagus and is nearly always fatal when diagnosed in the advanced stage. EAC develops through a slow stepwise progression through defined premalignant stages of disease, with microbiome involvement at each stage [5–7]. More specifically, EAC arises in a premalignant tissue called Barrett’s esophagus (BE), a relatively common condition in the US (1-2% prevalence with incidence continuing to rise) wherein the normal squamous epithelium of the distal esophagus near the gastroesophageal junction is replaced with specialized intestinal metaplasia [8]. BE is often the result of longstanding exposure to stomach acid as an adaptive response of the esophageal tissue in patients with symptoms of gastroesophageal reflux disease (GERD). In this way, a common pathway to progress to EAC from normal esophagus is through the stages of GERD then BE, and eventually EAC. Patients diagnosed with BE are recommended to undergo regular surveillance exams throughout their lifetime to look for potential early-stage EACs that are easier to treat than late-stage cancers [9]. Surveillance includes taking tissue biopsies, which allows a spatial and temporal map of EAC development to be studied. Although nearly all EAC develops from BE [8], the annual progression rate to EAC from BE is very low (<0.5% per year [10]) and thus most patients with BE undergo unnecessary surveillance. There is currently an unmet clinical need for improved cancer risk stratification in BE, including the development of reliable biomarkers for future EAC progression that enable the tailoring of cancer prevention strategies in a more personalized way. Therefore, EAC provides a useful system for developing temporal models, and by doing so, fulfills a critical knowledge gap in EAC prevention and the role of microbiome in cancer formation.

Recent studies have quantified the microbial communities within the esophagus at sequential disease states on the pathway to EAC [5,7,11–14]. These studies have shown shifts in the relative abundances of microbial taxa, Gram-negative versus Gram-positive bacteria, as well as microbial community diversity across stages of EAC progression. Additionally, characteristics of the oral microbiome have been shown to be associated with stages of progression in BE independent of patient oral health [15]. One dominant bacterium that has been examined extensively for its relationship to EAC is *Helicobacter pylori*. Although *H. pylori* is a known carcinogen with an established causal link to non-cardia gastric cancer and mucosa-associated lymphoid tissue (MALT) lymphoma [16], some epidemiological studies have demonstrated an inverse association with EAC [17–19]. A few studies even suggest the increase of widespread antibiotics and lower prevalence of *H. pylori* may be contributing to the 5-fold increase of EAC cases in the United States and other Western populations over the last few decades [6,20]. Although the absence of *H. pylori* has been shown to be associated with aneuploidy [21] and risk of advanced stages of high-grade dysplasia (HGD) and EAC in patients with BE [5], a mechanistic role of *H. pylori* in EAC progression, and the population dynamics of *H. pylori* growth in BE, have yet to be elucidated. Furthermore, a recent population-based study did not find an association with *H. pylori* eradication and EAC risk [22]. Overall, there is biological plausibility that altered microbiome and dysbiosis in BE may promote progression to EAC, but underlying mechanisms need to be defined. Here we investigate the microbial community dynamics evident in the stages of progression to EAC using mathematical models.

Existing microbiome studies of EAC relied on traditional statistical methods to test for differences in microbial species abundance between patient groups from single time-point data. These methods are often not designed for temporal dynamics and cannot be used to account for the expected variation in population sizes as the communities change over time. This requires time-dependent models of evolving populations during the progression to cancer. Relevant models of community assembly dynamics have been successfully applied to microbiome data of various environments such as compost, seawater, and sediment, as well as organisms such as roundworms, mice, and jellyfish [23]. They have also been applied to data on microbial communities within the human body such as comparing healthy versus diseased lung [24] and comparing human skin from differing geographical locations which found microbiomes from urbanized cities likely evolve through niche-based processes [25,26]. To similarly capture population dynamics of each microbial taxa in the community as well as continual migration of microbes into the esophagus, we developed models of community assembly specifically for EAC progression. By assuming certain growth and migration parameters for different species, we tested hypotheses such as the presence of predominantly neutral dynamics versus selective forces when fitting models to occurrence-abundance patterns measured in metagenomic data.

## Materials and methods

### Patient cohort information

Previous studies have compared the esophageal microbial communities in patients with normal esophageal tissue versus those with BE, and/or with EAC [11,12,14,27–31]. Few microbiome studies have separately categorized BE with low-grade and/or high-grade dysplasia [7,27], and we also do not examine this herein. One dataset we analyze in our study has the unique advantage of enabling the comparison of baseline whole genome sequencing data (WGS) data obtained from BE tissue samples in patients with BE who later progressed to EAC (‘cancer outcome’; CO) versus baseline WGS data from BE samples in patients with BE who did not progress to EAC (‘non-cancer outcome’; NCO) [32]. Eighty patients were included, each with two biopsy samples taken at two time points (320 total samples; see Supplementary Figure S1). The WGS data were obtained as part of a retrospective case-control study performed using the Seattle Barrett’s Esophagus Cohort (see [32] for details). The NCO patients remained cancer-free over median 17.47 years of active follow-up and across both time point 1 (TP-1) and time point 2 (TP-2). The CO patients had BE at (TP-1), and then progressed to EAC that was diagnosed (TP-2). For all patients at both TP-1 and TP-2, biopsy samples were taken at 2 different levels within their Barrett’s segment length (see [33] for exact distances from gastroesophageal junction and patient ages). Below we analyze the group of 80 samples from the 40 NCO patients at TP-1 (BE NCO cohort) versus the group of 80 samples from the 40 NCO patients at TP-1 (BE CO cohort) prior to their EAC diagnosis. Notably, all samples for this BE dataset were derived from DNA obtained from the isolated epithelium of the BE biopsies, rather than whole biopsy specimens.

To compare BE to the normal esophagus microbiome, we also analyzed WGS data obtained from esophageal brushings of normal esophageal tissue (n=50) as well as esophageal tissue from patients with gastroesophageal reflux disease (GERD; n=29). GERD is of interest because it is a high-risk condition for BE and symptomatic GERD (erosive esophagitis) is a common precursor stage. The samples were labeled as normal or GERD based on a histological report of the patient from which the sample was taken during an upper gastrointestinal endoscopy [14].

Additionally, we analyzed EAC (n=83) tissue samples from The International Cancer Genome Consortium (ICGC) to complete the healthy to cancer microbiome continuum (EAC cohort 1) [34]. Finally, to validate our findings we analyzed an additional EAC dataset (n=23) from patients who had visible BE (EAC cohort 2) [35]. Components of this dataset are now also included in ICGC and so, to ensure independence, we removed any overlapping samples from analysis of the ICGC dataset. A full description of samples analyzed in our study is provided in Supplementary Table S1.

### Computational host filtration pipeline and identification of microbial taxa

First, we remove human DNA from the WGS data using our validated computational pipeline (Supplementary Figure S2, see [36] for details). Briefly, host filtering is performed using minimap2 (v.2.17), mapping reads against the GRCh38.p14 and T2T-CHM13v2.0 human genome assemblies [37]. Remaining reads are considered non-host, and were mapped to known microbial species using the SHOGUN [38] pipeline and the Web of Life version 1 database [39]. To avoid false positive from potentially host-contaminated microbial reference genomes, we applied Exhaustive [40] and Conterminator [41] with human references GRCh38.p14 [42], T2T-CHM13v2.0 [43], and the Human Pangenome Research Consortium pangenomes [44] against the Web of Life, subsequently identifying 236 microbial taxa with regions of their genomes aligning with human reads (as was done with Web of Life-clean [40]). Out of an abundance of caution, we dropped any occurrences of the 236 potentially contaminated taxa from our taxonomy tables. In this pipeline, host filtering steps are essential to avoid the potential erroneous assignment of human DNA to microbial taxa. We note that the procedure outlined above was performed across all datasets analyzed in main results (Supplemental Table 1) and for 377 buccal mucosa samples from the Human Microbiome Project that were used to evaluate *Helicobacter pylori* prevalence in oral communities [45,46].

To reduce risk of false positives from short read multimapping, we applied a novel coverage dispersion filter called *micov* which eliminated taxa with low genomic coverage (https://github.com/biocore/micov). Using micov, we determined coverage dispersion thresholds for microbial genomes that were represented or mismapped by testing on a *Staphylococcus aureus* monoculture [47] subset to various sequencing depths. We were able to distinguish *Staphylococcus aureus* and other similar species in the *Staphylococcus* genus from all other microbial mis-mappings with a sensitivity of 87.8% and specificity of 95.18%. Our criteria for determining if DNA reads were mis-mapped to a particular taxon when genome coverage was less than 10% across all samples (those taxa with >10% coverage kept, as determined using Zebra [47]) were: A) there were at least 4 unique areas of the taxa with mapped reads, B) greater than 75% of reads mapped to the same 3 consecutive bins of the genome, or C) greater than 75% of reads mapping to the taxon are only mapped to a single consecutive region of the genome in any one sample (see Supplementary Methods for details). Taxa meeting any of the 3 criteria above were removed from downstream analyses. For the BE NCO and BE CO dataset, only criteria A was used for filtering due to the low microbial biomass of these samples. Within each dataset, samples were rarefied and samples with less than the rarefaction depth were removed (Supplementary Figure S3). All taxa were collapsed to species level unless otherwise stated.

Additionally, to confirm our modeling results were robust to varying microbial DNA classification approaches, we also analyzed the BE NCO and BE CO datasets using MetaPhlAn4 with the flags “-t rel_ab_w_read_stats --unclassified_estimation” (run using the Nextflow workflow available at https://github.com/FredHutch/metaphlan-nf commit 4a31444, and the reference database version mpa_vJan21_CHOCOPhlAnSGB_202103) [48]. Notably, MetaPhlAn was selected as a top performing tool in the Critical Assessment of Metagenome Interpretation Round I [49] and Round II [50]. Finally, we performed an additional analysis that restricted mapping exclusively to a list of previously documented human-associated microbes (see Supplementary Table S1A from [3]).

### Statistical analysis of metagenomic data

All statistical analysis of the metagenomic abundances generated using the methods described above were performed using QIIME2 (v. 2023.2.0) [51] using the API in Python (v. 3.8.16). Beta-diversity was calculated using dimensionality reduction of the BIOM table with Gemelli’s Robust Principal Component Analysis (RPCA) function using DEICODE (v. 0.2.4) [51,52] to create a distance matrix for PERMANOVA and RPCA-Principal Component Analysis plots.

### Community assembly modeling of human esophageal microbiome

Figure 1A illustrates the model setup. The esophageal microbiome contains different microbial species undergoing growth dynamics described mathematically by stochastic birth-death-immigration processes. We assume there are approximately constant *N* microbial cells in an adult patient’s esophageal microbiome community and *N*_*i*_ represents the number of microbes of species *i* in the esophagus (thus *N = ΣN*_*i*_). When an individual microbial cell dies in the esophagus, it can be replaced by either another microbial cell that replicates in the present esophageal community or it can be replaced by a microbial cell entering the microenvironment from a source pool. Immigration of microbes from a source pool to the esophagus likely includes microbes from the oral cavity, proximal stomach, and the adjacent normal tissue in the case of BE microbiome. Immigration allows new microbes to enter the esophagus eliminating the unrealistic fixation of a single microbial species with the above dynamics. Following the mathematical theory developed previously for these modeling assumptions [23,53], we applied two different approaches for building the community assembly models based on the presence/absence of selection of certain microbial species within the community.

**Figure 1.**
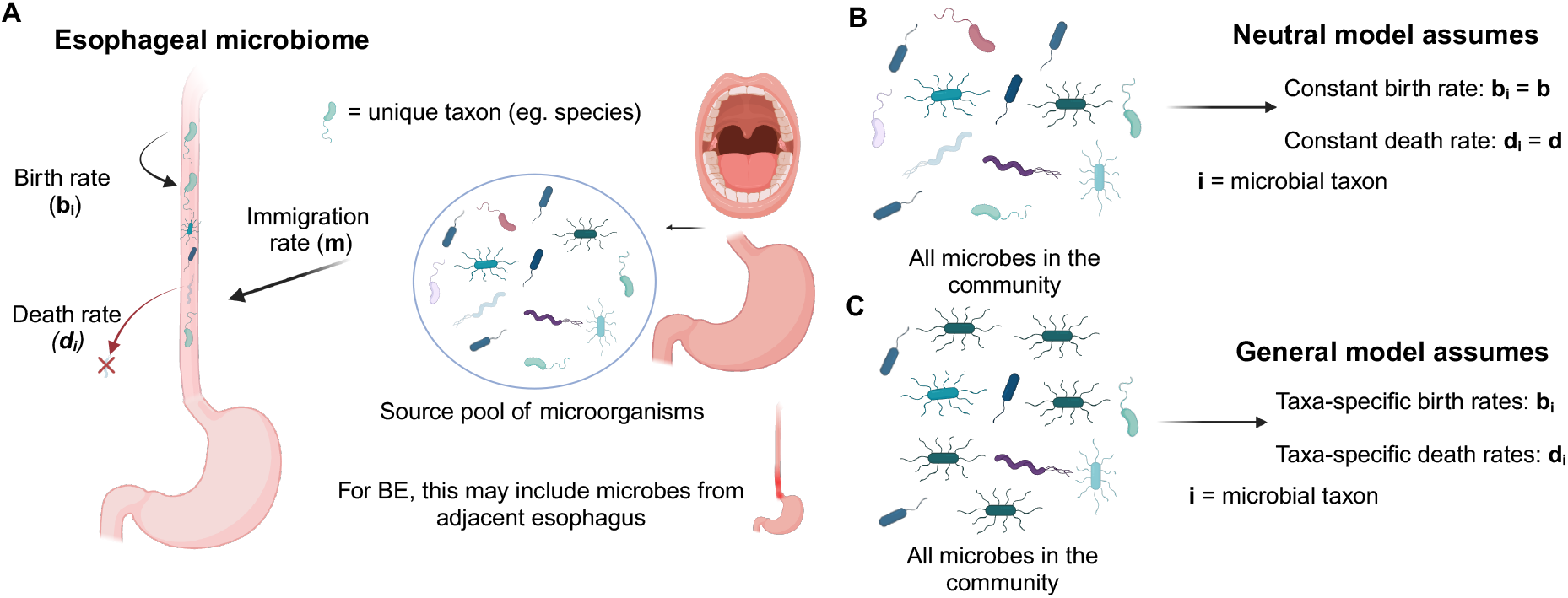
Neutral and non-neutral community assembly and model dynamics in the context of the esophagus microbiome. **A**. An overview of the model. **B**. Neutral model with mixed microbes in the community and an equal birth and death rates among all microbial taxa. **C**. Non-neutral model with potential domination of particular microbial taxa in the community enabled by different birth and death rates among microbial taxa.

The first approach is built on neutral theory [54] and thus assumes the rates for birth (*b*_*i*_) and death (*d*_*i*_) are the same for all microbes in a community (i.e., *b*_*i*_ *=b* and *d*_*i*_ *=d*). Thus in this model, no microbial species has a selective growth advantage over any other (Figure 1B). Here, we are defining a selective advantage as an increased birth or decreased death rate for one microbe over others. The second more general approach is built on a non-neutral model called niche theory which assumes that particular microbes may have a selective advantage over others. This model accounts for differences in proliferation and death rates across microbial species, allowing for certain microbes to differ in abundance and probability of extinction (Figure 1C).

When a single microorganism dies, the probability of being replaced by a microbe outside the present esophageal community is represented by *m*, known as the immigration rate. Alternatively, the probability of being replaced by a species present in the esophagus is *(1-m)*. The probability of a particular species of microbe being chosen for replacement with a birth event is based on the abundance of that microbe in the esophagus *(N*_*i*_ */N)* and the source community pool probability for that microbe (*p*_*i*_). As discussed in Sieber et al. [23], we can then calculate the probability that the abundance of species *i* may increase, decrease, or remain unchanged following the death of a microbe in the local microorganism community using the transition probability master equations corresponding to Hubbell’s original neutral model [54].

If *N* is large, we can then assume the relative abundance of species *i* in the esophagus, *x*_*i*_ *= N*_*i*_ */N*, to be approximately continuous and use the Fokker-Planck equation to model the continuous approximation for the probability density function *Φ_i_(x_i_; N, p_i_, m)*. We can then approximate the long-term equilibrium solution of this equation by (see [23] for full derivation).

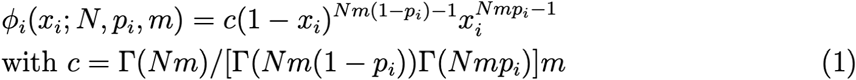

To connect this to empirical data, we use nonlinear least squares minimization to fit the observed occurrence frequencies f_i_ from the rarefied data to the following truncated cumulative probability density function for observing a particular species in a local community/sample,

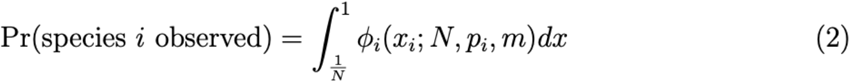

where *N* is the total number of species counts per sample and the relative abundance of species *i* in the source pool (*p*_*i*_) is approximated by the mean relative abundance of species *i* across all samples in a given dataset. Thus with *p*_*i*_ and the total microbial population in the esophagus (*N*) defined, the only free parameter to fit to the data is the immigration rate *m*. Additionally, we calculated the standard coefficient of determination (*R*^*2*^*=1 -* the ratio of the sum of squared residuals and the total sum of squares) to provide a measure of goodness of fit for best-fit neutral models in each dataset (see ref [23] for details).

### Incorporation of selection into model structure and Gillespie simulations

In the non-neutral model, we assume that each microbial species *i* has a species-specific birth rate (*b*_*i*_) and death rate (*d*_*i*_) that can vary from one another. We assume that the same evolutionary dynamics for immigration apply as above-when a microbe dies, it can either be replaced by a microbe inside the source pool or resident within the esophagus. In the non-neutral case, the probability of a microbe being chosen for replacement is not based solely on the relative abundance of the microbe but also on birth and death rates, so we modify the transition rates to take into account both the microbial taxa member that dies (*j*) and the microbial taxa which will increase in number (*i*) when calculating the abundance of microbes in the esophagus (*N*).

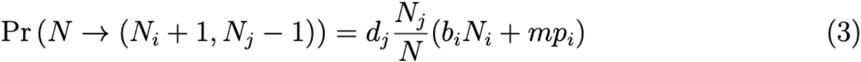

In this way, the transition probabilities were modified to account for both the increase of species *i*, and the decrease of species *j*.

The probability distribution for the community composition in the non-neutral system can be analytically solved at equilibrium for the case of two different microbial species. However, in a realistic model for a local community considering potentially hundreds of microbial species identified in human tissue samples, the size of the system consisting of transition probabilities becomes too large to be computationally feasible, as discussed in Zapién-Campos et al. [53]. Therefore, we use the Gillespie algorithm to perform stochastic simulations of this model using transition probabilities in Eq. (3) in order to compute the mean relative abundances and occurrence frequencies of the evolving microbial populations needed to characterize the evolutionary dynamics [53]. When performing simulations, we used the number of hosts, taxa, and microbes per host derived from the dataset of interest with a fixed time simulation of t=10. To set the initial conditions of the community assembly for each sample in the simulated data at a starting time point, we randomly drew *N* microbes according to the source pool distributions p_i_, where we again approximate this distribution by using the microbial relative abundances from the taxonomic table for that dataset. We are then able to test different hypotheses on selective growth advantages for certain microbial species and compare simulated results to real data.

## Results

### Microbial diversity differs across natural history of EAC progression

We first quantified the microbial species present in each tissue sample by mapping DNA sequencing reads to curated databases of human and microbial genomes (see Materials and methods), and then compared relative abundances between esophageal datasets (Supplementary Figure S4). We found similarities at the phylum level across disease states, although healthy esophagus and GERD had higher levels of Firmicutes and Bacteroidetes compared to others. We found many shared taxa between datasets at the species level (Supplementary Figure S5). Alpha-diversity was measured across each disease-stage group using both richness (Figure 2A) and Shannon entropy (Figure 2B). The median Shannon entropy metrics across disease states were: Healthy 4.82 (IQR=4.54-5.21), GERD 5.17 (IQR=4.64-5.38), BE NCO 3.79 (IQR=3.71-3.98), BE CO 3.76 (IQR=3.67-3.85), EAC cohort 1 2.48(IQR=2.26-2.81), EAC cohort 2 2.90 (IQR=2.84-2.92). We found statistically significant differences across all groups in alpha-diversity (Kruskal-Wallis p=4.81 × 10^−43^; Shannon entropy) and then in post-hoc Dunn’s tests pairwise comparisons between disease stages of Healthy/GERD vs. BE vs. EAC (p<0.01, Shannon entropy). Non-significant differences in pairwise comparison Dunn’s tests (Shannon entropy) were found for Healthy versus GERD, BE CO versus BE NCO, and EAC cohort 1 versus EAC cohort 2. For healthy/GERD samples, we found a high richness (median = 296 in healthy esophagus, 309.5 in GERD) and largest Shannon entropy metrics. This is potentially influenced by the use of esophageal brushings rather than esophageal biopsies to obtain these samples. Overall, we found decreasing Shannon entropy as stages progressed to EAC, which is consistent with previous findings [11,12].

**Figure 2.**
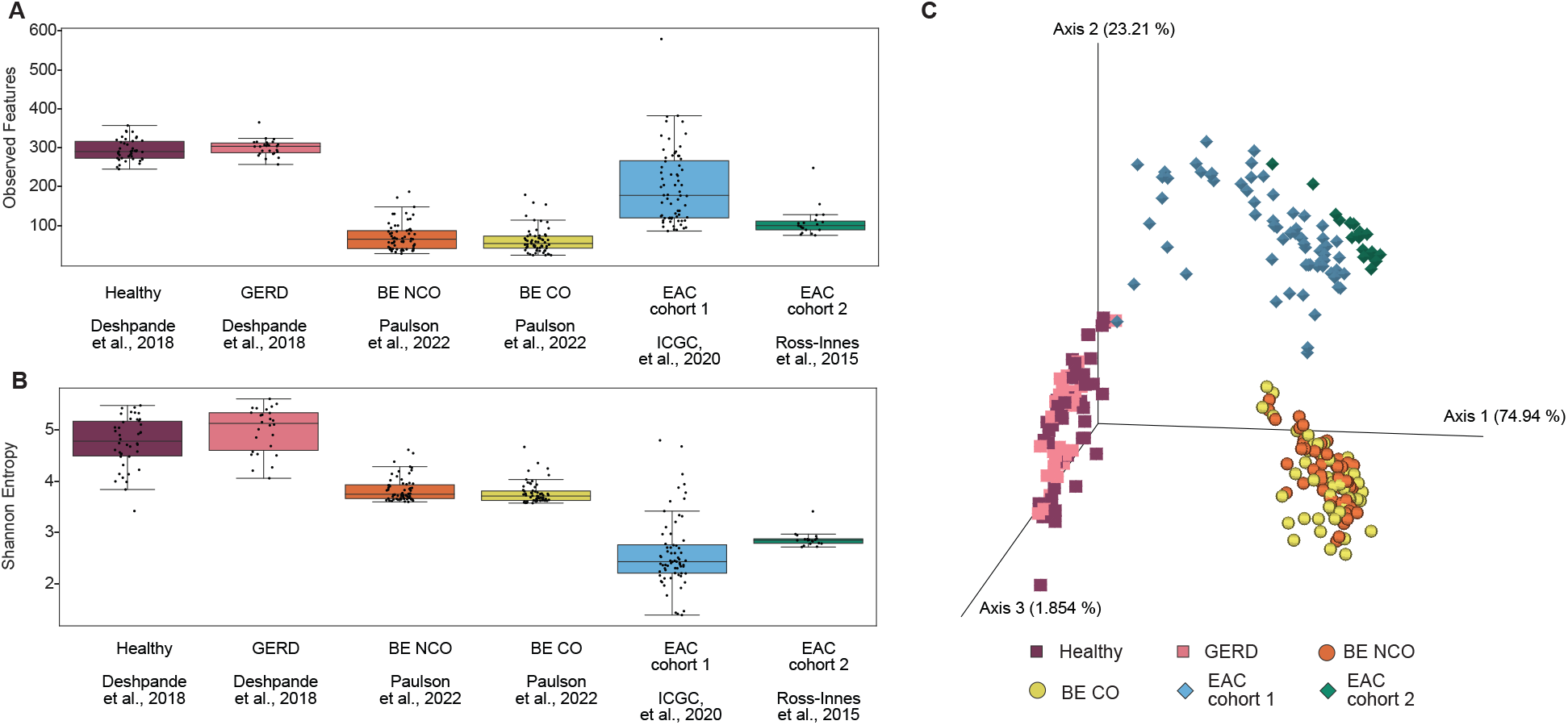
Alpha- and beta-diversity metrics across disease sets. **A**. Microbial alpha-diversity represented by taxon richness (i.e., unique observed features) across disease stages. **B**. Microbial alpha-diversity represented by Shannon entropy across disease stages. **C**. Differences in community structure represented by the RPCA-PCoA between disease stages (PERMANOVA *p*=0.0001, pseudo-*F*=487). Shape corresponds to the disease state: square = Deshpande et al., 2018, circle = Paulson et al., 2022, diamond = Ross-Innes et al., 2015 & International Cancer Genome Consortium (ICGC) et al., 2020.

We also calculated beta-diversity with DEICODE [52] using a Robust principal component analysis (RPCA; Figure 2C). We observed clustering among disease state groups: Normal and GERD samples, BE samples, and EAC samples. As with alpha-diversity, we observed using pairwise PERMANOVA (pseudo-*F* statistic) that all datasets were statistically significantly different (p=0.0001). This holds across all pairwise comparisons (p=0.0001) except for healthy versus GERD (p=0.4801) and BE NCO versus BE CO (p=0.5381).

### Neutral models generate occurrence-abundance patterns seen in development from normal esophagus to BE

Although comparing exact taxonomic differences between datasets may be influenced by sample collection methods and thus the microenvironments included in the sequencing reads (see Supplementary Table S1), comparing population dynamics predicted by community assembly models can better avoid such biases because conclusions are not based as strongly on specific taxa. We first applied the neutral model (where no bacterial species has a growth advantage over another) to all esophagus disease-stage subsets (Figure 3, see Materials and methods). In general, we found high *R*^2^ values for samples from normal esophagus and GERD (*R*^2^>0.80) and BE (*R*^2^>0.65) suggesting that neutral dynamics can explain the majority of species populations in these cohorts. We found occurrence-abundance patterns measured in BE NCO and CO samples were consistent across time points, evidence that the total community structure is likely at steady state (Supplementary Figure S6).

**Figure 3.**
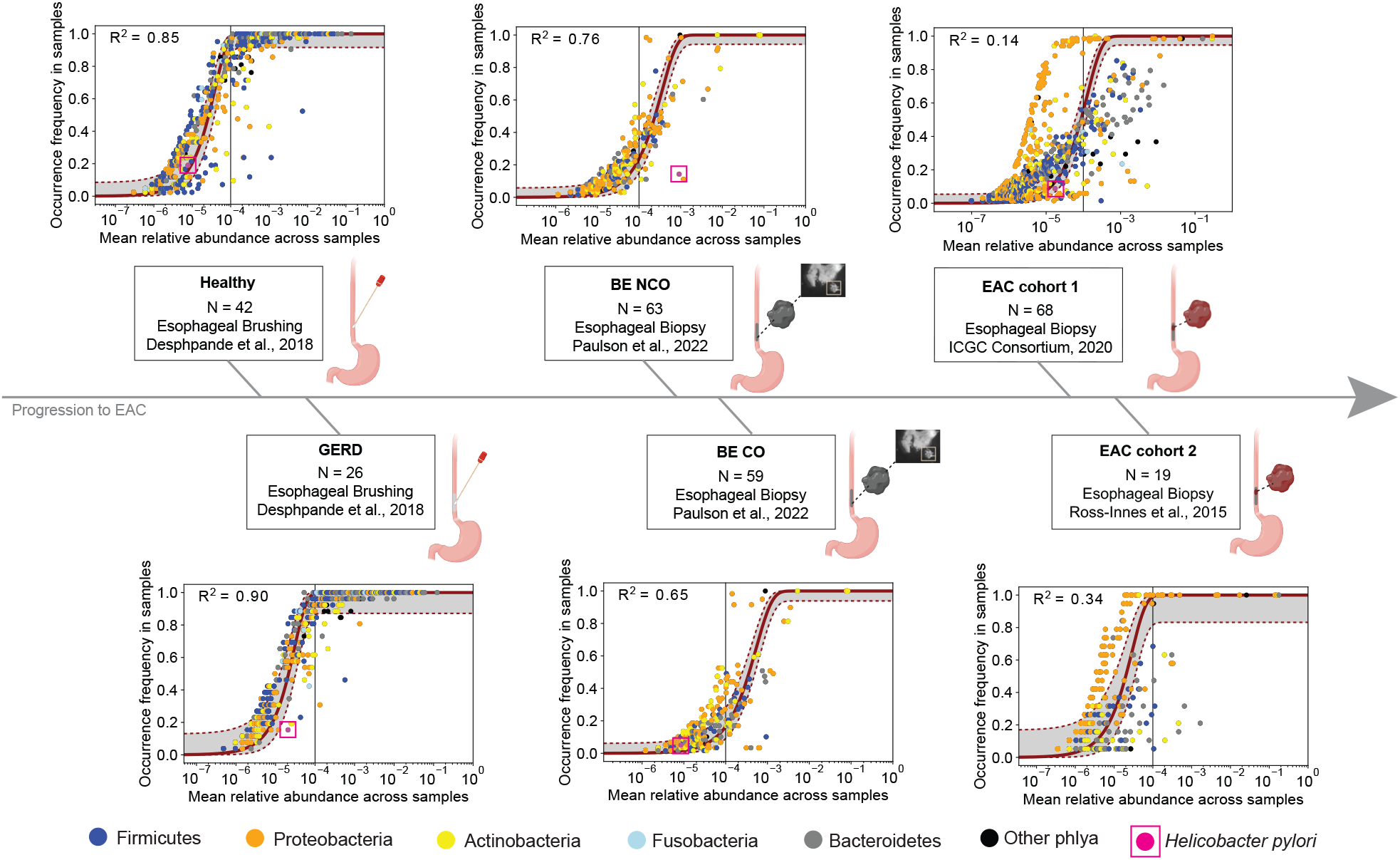
Community assembly dynamics in progression to EAC. Plots are ordered in sequence of disease state progression from healthy to EAC (see Supplementary Table S1 for patient/sample inclusion). The red curves represent the neutral model fit to data points representing unique taxa at the species level. The gray regions delineated by red dotted lines represent 95% bootstrap confidence intervals (obtained by resampling the hosts 100 times with replacement and refitting) and the *R*^2^ value represents goodness of fit to the neutral model. The color of the data point represents phyla. *Helicobacter pylori* is colored pink with a box outline. Vertical line indicates mean relative abundance = 10^−4^. *H. pylori* is found with the following occurrence frequency in each group: Healthy 19.0476%, GERD 15.3846%, BE NCO 14.2857%, BE CO 5.0847%, EAC cohort 1 8.8235%, EAC cohort 2 0%.

Neutral model goodness-of-fit *R*^2^ values decreased as stages progressed from normal to EAC. In the case of EAC cohort 1, the neutral model provides a poor fit to the data (*R*^2^=0.14), suggesting potential evidence for non-neutral microbial dynamics in EAC patients. In particular, we see there may be some species undergoing more neutral dynamics (S-shaped curve of data points for EAC), but a large number of microbial species are found in higher abundance than expected by the neutral expectation. The EAC samples had the largest number of microbial taxa deviating from the neutral curve fit. Similar results for the EAC occurrence-abundance pattern were found in the Ross-Innes et al. EAC cohort 2 dataset (*R*^2^=0.34, Figure 3).

In Figure 3, the top five most common phyla found in the esophagus are denoted by different colors and *H. pylori*, which has been found to colonize distal esophagus and aggravate reflux injury [55,56], is shown in pink in each disease-stage panel. As noted above, *H. pylori* is of particular interest because of its inverse association with BE [19] and EAC development [57] in some studies, yet causal association in non-cardia gastric adenocarcinoma through an analogous intestinal metaplasia-dysplasia-adenocarcinoma sequence (also known as the Correa cascade) [16]. We identified *H. pylori* in samples from every disease stage except for EAC cohort 2 samples from Ross-Innes et al [35]. The only instances that data for *H. pylori* deviates significantly from the neutral curve fit was in the GERD group and the BE NCO group. This result holds when sub-setting the cohorts by sex as an EAC risk factor (males have higher risk of EAC than females [58]), except in the case of males with GERD where occurrence-abundance of *H. pylori* was close to the neutral expectation. Although the sample size for the male GERD patient subset was modest (n=10), we also found *H. pylori* at a lower mean relative abundance (1×10^−6^) compared with female GERD patients (n=19) who had a mean relative abundance of 3.4×10^−5^.

To validate these findings, we fit the neutral model to BE NCO and CO datasets after preprocessing with the MetaPhlan4 pipeline (see Materials and methods). We identified fewer microbial species overall with this method, but *H. pylori* is still identified as likely neutral in only the BE CO cases (Supplementary Figure S7). Finally, we restricted our microbial taxa to those previously documented as human-associated and still found that 1) normal esophagus, GERD and BE show evidence for neutral dynamics while EAC does not, and 2) GERD and BE NCO are the only cases where *H. pylori* is outside the 95% confidence interval for expected neutral dynamics (Supplemental Figure S8). The top 10 bacterial species with largest deviation from the 95% confidence intervals for a neutral model fit found in each of the BE NCO and CO datasets are provided in Supplementary Tables S2 and S3. Other non-neutral microbes beyond *H. pylori* included *Klebsiella pneumoniae* in the NCO BE dataset (nearly identical occurrence-abundance as *H. pylori*), *Prevotella copri* in the BE CO cohort, and *Neisseria mucosa* in both cohorts.

### *Helicobacter pylori* exhibits positive selection in non-progressing BE

Because *H. pylori* specifically deviated from the neutral model fit in the BE NCO cohort (Figure 3), we performed Gillespie simulations for neutral and non-neutral scenarios for this dataset. This allowed us to use the relative abundances of taxa observed in the BE NCO tissue samples but vary the birth and death rates of each microbe to evaluate what assumptions for dynamics would represent the data most closely. In the BE NCO neutral model simulation, we found that *H. pylori* was predicted to have occurrence-abundance data close to the neutral fit curve (Supplemental Figure S9A) unlike the actual data (Figure 4A). Additionally, simulated *H. pylori* occurrence frequency was much higher than the occurrence frequency measured in the actual data when assuming a source pool prevalence for *H. pylori* equaling its mean relative abundance in the BE data (Supplemental Figure S9A). In Supplemental Figure S9B, we show different simulation results for *H. pylori* with varying birth and death rates and source pool conditions. We found consistent data on *H. pylori* occurrence-abundance in simulations compared to the empirical data (pink dot with box outline) when assuming a source pool prevalence probability of 1e-05. To examine this assumption in independent data, we found that *H. pylori* was present in 39 of a total of 377 (15%) buccal mucosa samples from the Human Microbiome Project [45,46] at a mean relative abundance of 1.13e-05 [45]. Thus, with an assumption that the oral microbiome mainly contributes to the source pool for BE microbial communities, we used 1e-05 for *H. pylori* source abundance for all further model simulations.

**Figure 4.**
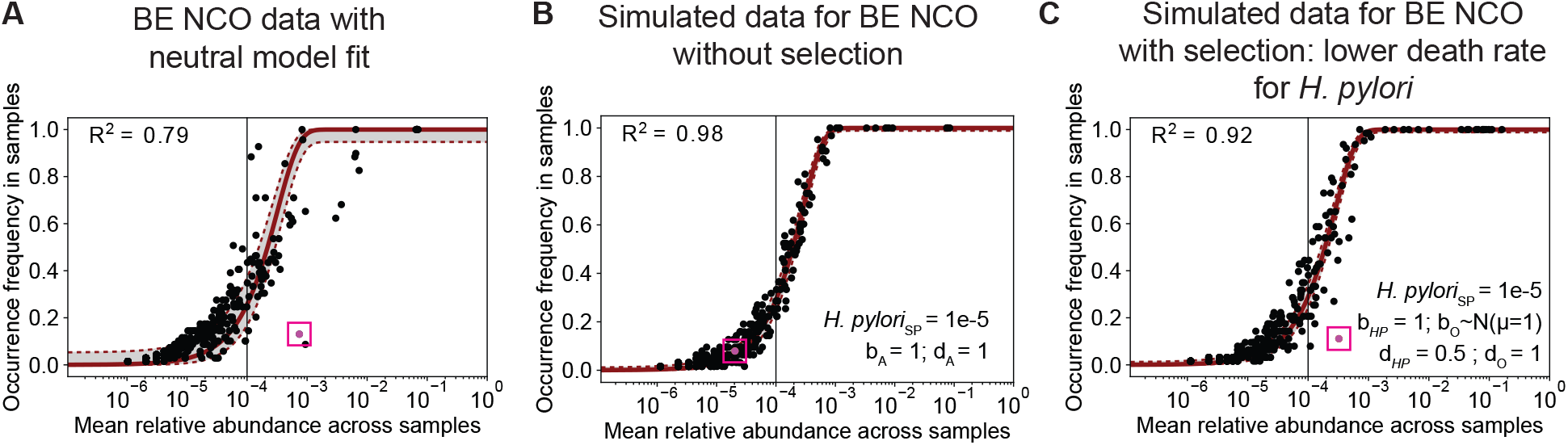
Simulation results for BE patients with non-cancer outcome (NCO). **A**. Same data as shown for BE NCO panel in Figure 3, with neutral model fit for BE NCO patients. **B**. Neutral simulation with data from the BE NCO patients using the adjusted source pool conditions for *H. pylori*. All (‘A’) microbes have an equal birth and death rate of 1. **C**. Non-neutral simulation for BE NCO data with *H. pylori* adjusted source pool prevalence (*H. pylori*_SP_ *=*1e-5). For the *H. pylori* taxon, the birth and death rates are 1 and 0.5, respectively. For all other microbes, birth rates are drawn independently from a normal distribution (mean=1, standard deviation=0.1) and the death rate is set equal to 1. For **A-C**, the pink dot depicts data for *H. pylori*. For **B-C**, the source pool conditions of the *H. pylori* taxa are set to 1e-5 (see Results and Supplemental Figure S9 for details). SP = source pool; HP = *H. pylori*.

Even with a neutral model fit with this source pool assumption, simulated *H. pylori* occurrence-abundance does not recapitulate the data (Figure 4A,B). Therefore, to test the effects of potential non-neutral dynamics, we modified the simulations by assuming that *H. pylori* has a selective advantage over other microbials species. We tested a variety of different parameter settings to compare to the data in Figure 4A and found that assuming a lower death rate for *H. pylori* (d_HP_=0.5) compared with other microbes (d_A_=1) accurately reflected the actual data (Figure 4C). We also experimented with variable birth rate and found increased birth rate gives a similar result and fit to the data (Supplemental Figure S9C). Overall, higher expected growth rates (birth - death) for *H. pylori* align with their known predominance in bacterial populations of the stomach when present.

### Neutral dynamics do not explain occurrence-abundance patterns in EAC tissue

Because we observed that the neutral model did not fit well to the data for EAC samples, we also performed stochastic simulations based on this dataset to test effects of differing assumptions for microbial population dynamics. With relative abundances from the EAC cohort 1 used as the source pool prevalence for each species, we ran neutral model simulations where the birth and death rates across all microbes were set equal to 1 and resulting occurrence-abundance did not reflect the data, as anticipated (Figure 5A,B). We next created simulated data with a random birth rate ranging from 1 to 6 for each microbial species and a constant death rate equal to 1 for each species, and found *R*^2^ = 0.59, which more closely resembles the empirical data (Figure 5C). We re-ran the simulations again with a random death rate ranging from 1.5 to 4 for each microbe and a constant birth rate equal to 4 for each species, and found a similar pattern to the original data with *R*^2^ = 0.74 (Supplemental Figure S10A). Finally, we varied the birth rate (normally distributed with mean=1, standard deviation = 0.1) as well as varied the death rate (normally distributed with mean=1, standard deviation = 0.1) and found that these models did not recapitulate the original data (Supplementary Figure S10 B,C).

**Figure 5.**
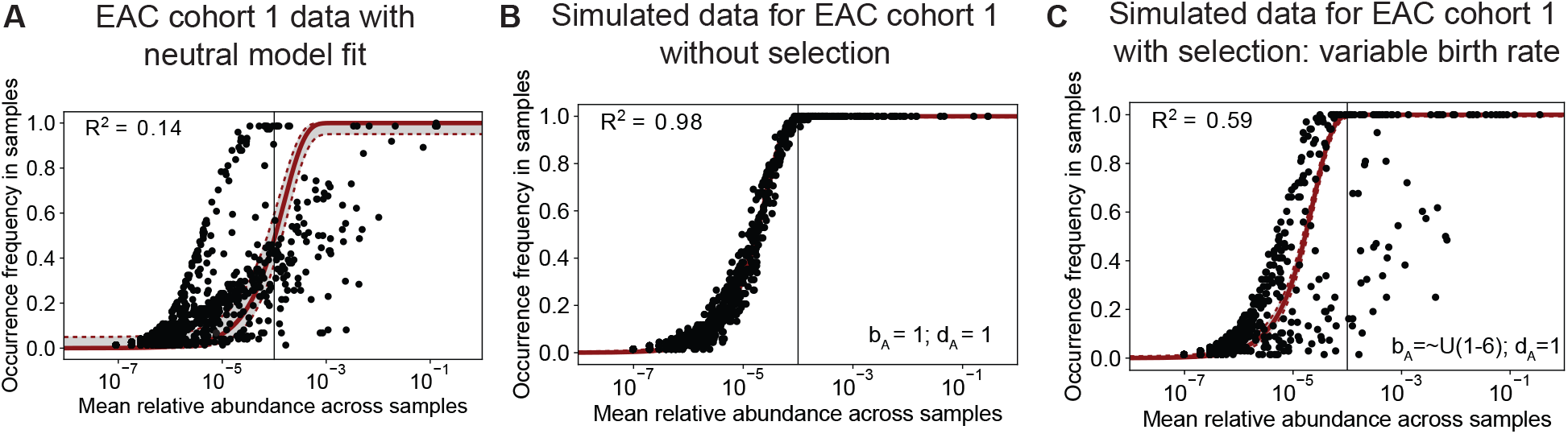
ICGC EAC cohort 1 data with fits assuming neutral model at steady-state, neutral simulation, and non-neutral simulation. **A**. Same data as shown in Figure 3, neutral model fit to data from EAC cohort 1 patients in ICGC dataset. **B**. Neutral model simulation with data from the EAC cohort 1 patients in ICGC dataset. All (‘A’) microbes have an equal birth and death rate of 1. **C**. Non-neutral model simulation with all microbes having variable birth rate drawn independently from a uniform distribution ranging from 1 to 6, and a death rate of 1. ICGC = International Cancer Genome Consortium.

## Discussion

With increasing cases of both BE and EAC in Western populations, new approaches to better understand contributions to EAC risk are essential. As was found with the established role of *H. pylori* in the formation of peptic ulcers and gastric cancers, and the role of a diverse oral microbiome in oral and systemic health [59], the esophagus also provides a microenvironment for microbial populations to thrive and potentially contribute to patient health and disease. The communities have been quantified previously in numerous studies, mainly using 16S rRNA sequencing, and may influence progression to BE and subsequently EAC but the expected population dynamics of these communities has not been previously explored. Therefore, our aim was to better understand the communities within the esophageal microbiome as a patient progresses from normal tissue to EAC.

To explore dynamics within the esophageal microbiome during EAC progression, we utilized whole genome sequencing (WGS) data from esophageal tissue samples and applied a rigorous host depletion pipeline to ensure we removed all human DNA reads [36] and mapped any potentially microbial reads to an extensively cleaned database (WOL-clean) as well as using a coverage filter called micov. Comparing histological subtypes at each disease stage using the ecological diversity measure of Shannon entropy, we found decreasing diversity from normal esophagus to GERD to BE to EAC. Study limitations in these analyses include varying sampling techniques and centers (Supplemental Table S1), as well as a lack of reagent ‘blank’ controls and positive spiked controls run in parallel, which would increase confidence in minimizing contamination [53]. Additionally, we did not assess differences between more advanced premalignant stages within BE such as low-grade dysplasia and high-grade dysplasia.

The effects on our results introduced by these limitations are minimized when considering the community dynamics with rarefied data, where we independently assess data for each disease stage and compare observed occurrence-abundance patterns across stages of progression. We observed that patterns in earlier stages of progression (normal esophagus, GERD, and BE) can be recapitulated with assumption of a neutral model. Zapien-Campus et al. note that such patterns can in fact be recreated by specific assumptions for non-neutral dynamics [60] but the hypothesis of neutrality cannot be rejected for this data. In contrast, patterns in the data for EACs appear quite different, and only a subset of the microbial taxa appear to fit a neutral expectation in their occurrence-abundance patterns. This may imply that more of the EAC microbial populations are undergoing selection pressures in the tumor microenvironment. By simulating data using mathematical models of microbial evolution, we can quantify the parameter regimes of differential selection (as measured by population growth rates) that most closely match the empirical data.

We specifically examined *H. pylori* due to its potentially protective role in EAC development. Across disease states, the mean occurrence frequency of *H. pylori* was ∼10%, however, its mean relative abundance across samples was greatest in the NCO group (10^−3^) compared to other groups (<10^−4^). This may suggest that *H. pylori* has an active role in preventing progression to EAC in NCO patients and BE in patients with GERD, or at least delaying the onset, but further mechanistic studies are needed, particularly since confounding cannot be ruled out here. The only other group that demonstrated non-neutral *H. pylori* was the GERD group. These patients have not yet crossed over the critical threshold to BE and further stages but may develop BE at a later time point.

Additionally, other microbes in the NCO BE dataset were found to be under possible selection including *Klebsiella pneumoniae* which had a very similar abundance and occurrence to *H. pylori. K. pneumoniae* is often found in the human gut but causes serious infections across other tissue types [61] and was linked to Acute Esophageal Necrosis in a case study [62]. There were a few disease-specific microbes that were identified as non-neutral in both cohorts, including *Neisseria mucosa*, commonly found in the oral and nasal mucosa but can cause upper respiratory infection [63]. *Prevotella copri* was only found to be non-neutral in the BE CO cohort, and is often found in the human gastrointestinal microbiome with varying positive and negative effects [64], but studies have found correlations between *P. copri* abundance and exacerbated intestinal mucositis [65] as well as colitis [66] in mice. Additionally, *P. copri*, similar to *H. pylori*, is implicated in gastric carcinogenesis [67] and was associated with an increased risk for gastric cancer development in a Korean population [67]. Finally, when we modeled the assumption that *H. pylori* has a higher survival rate compared to other microbes in the esophagus, which was motivated by the observation that *H. pylori* unequivocally dominates the gastric microbiome when present, the data recapitulated what we observe in BE non-progressors. One hypothesis is that the dominance of *H. pylori* in the esophagus is protecting against the establishment of microbes that are detrimental to the esophagus and promote BE transformation to EAC. Certainly, studies evaluating the impact of *H. pylori* virulence factors (e.g., CagA), *H. pylori* eradication, as well as the impact of gastric histology (e.g., gastric atrophy with or without metaplasia) and gastric pH, factors both influenced by *H. pylori*, on development and progression of BE would be informative. Although further validation is needed, our results suggest microbes such as *H. pylori* in precursor stages of GERD and indolent BE, as well as substantial numbers of taxa in the EAC tissue microenvironment, likely undergo selective pressures that influence relative abundances measured throughout carcinogenesis.

## Conclusion

In summary, there is evidence for neutral population dynamics for the majority of microbial species in normal esophagus, GERD, and Barrett’s esophagus tissue microenvironments. This implies that fluctuations in relative abundances measured in previous studies at various time points and disease stages may simply reflect such underlying dynamics. Our data also suggest that *Helicobacter pylori* in particular may be under positive selection in pre-cancer stages, and EAC microenvironments exhibit population dynamics that are largely non-neutral. Bacteria are present in precancerous stages of esophageal adenocarcinoma, and rigorous bioinformatics methods and mathematical models are needed to identify these communities along with their evolutionary dynamics from WGS data in order to better understand what influence they may have on cancer progression.

## Data availability

All data used in our study has been previously published [14,32,34,35], as outlined in Supplementary Table S1.

## Code availability

Code to recreate all results and apply methods to other datasets can be found on GitHub: https://github.com/cguccione/comad_of_EAC_progression. The code created for modeling analyses was adapted from [53] and [53]. The workflow used to run MetaPhlAn4 is available at https://github.com/FredHutch/metaphlan-nf.

## Acknowledgments

This work was supported by AGA Research Foundation (AGA Research Scholar Award AGA2022-13-05) and NIH R01 CA270235 to K.C. The study was supported in part by the NIDDK-funded San Diego Digestive Diseases Research Center (P30 DK120515). Additionally, this work was supported by NIH grants (R01 CA241728, P30 CA023100, NIH/NIGMS T32GM007198, NCI U24CA248454) to R.K. J.P.S. is supported by grants SD IRACDA, Professors of the Future (5K12GM068524-17) and USDA NIFA (2019-67013-29137). T.P. is supported by NIH grants (R01 CA140657, R01 CA270235 and R21 CA259687). S.C.S. receives funding from a Veterans Affairs Career Development Award (ICX002027A). Figure 1 was made with BioRender.

## References

1. Nejman D, Livyatan I, Fuks G, Gavert N, Zwang Y, Geller LT, et al. The human tumor microbiome is composed of tumor type-specific intracellular bacteria. Science. 2020;368: 973–980.

2. Dohlman AB, Klug J, Mesko M, Gao IH, Lipkin SM, Shen X, et al. A pan-cancer mycobiome analysis reveals fungal involvement in gastrointestinal and lung tumors. Cell. 2022;185: 3807–3822.e12.

3. Battaglia TW, Mimpen IL, Traets JJH, van Hoeck A, Zeverijn LJ, Geurts BS, et al. A pan-cancer analysis of the microbiome in metastatic cancer. Cell. 2024;187: 2324–2335.e19.

4. Sepich-Poore GD, Guccione C, Laplane L, Pradeu T, Curtius K, Knight R. Cancer’s second genome: Microbial cancer diagnostics and redefining clonal evolution as a multispecies process. Bioessays. 2022;44: e2100252.

5. Guccione C, Yadlapati R, Shah S, Knight R, Curtius K. Challenges in determining the role of microbiome evolution in Barrett’s esophagus and progression to esophageal adenocarcinoma. Microorganisms. 2021;9: 2003.

6. Snider EJ, Freedberg DE, Abrams JA. Potential Role of the Microbiome in Barrett’s Esophagus and Esophageal Adenocarcinoma. Dig Dis Sci. 2016;61: 2217–2225.

7. Snider EJ, Compres G, Freedberg DE, Khiabanian H, Nobel YR, Stump S, et al. Alterations to the Esophageal Microbiome Associated with Progression from Barrett’s Esophagus to Esophageal Adenocarcinoma. Cancer Epidemiol Biomarkers Prev. 2019;28: 1687–1693.

8. Curtius K, Rubenstein JH, Chak A, Inadomi JM. Computational modelling suggests that Barrett’s oesophagus may be the precursor of all oesophageal adenocarcinomas. Gut. 2020;70: 1435–1440.

9. Shaheen NJ, Falk GW, Iyer PG, Souza RF, Yadlapati RH, Sauer BG, et al. Diagnosis and Management of Barrett’s Esophagus: An Updated ACG Guideline. Am J Gastroenterol. 2022;117: 559–587.

10. Desai TK, Krishnan K, Samala N, Singh J, Cluley J, Perla S, et al. The incidence of oesophageal adenocarcinoma in non-dysplastic Barrett’s oesophagus: a meta-analysis. Gut. 2012;61: 970–976.

11. Elliott DRF, Walker AW, O’Donovan M, Parkhill J, Fitzgerald RC. A non-endoscopic device to sample the oesophageal microbiota: a case-control study. Lancet Gastroenterol Hepatol. 2017;2: 32–42.

12. Radani N, Metwaly A, Reitmeier S, Baumeister T, Ingermann J, Horstmann J, et al. Analysis of Fecal, Salivary, and Tissue Microbiome in Barrett’s Esophagus, Dysplasia, and Esophageal Adenocarcinoma. Gastro Hep Adv. 2022;1: 755–766.

13. Yang L, Lu X, Nossa CW, Francois F, Peek RM, Pei Z. Inflammation and intestinal metaplasia of the distal esophagus are associated with alterations in the microbiome. Gastroenterology. 2009;137: 588–597.

14. Deshpande NP, Riordan SM, Castaño-Rodríguez N, Wilkins MR, Kaakoush NO. Signatures within the esophageal microbiome are associated with host genetics, age, and disease. Microbiome. 2018;6: 227.

15. Solfisburg QS, Baldini F, Baldwin-Hunter BL, Lee HH, Park H, Freedberg DE, et al. The Salivary Microbiome and Predicted Metabolite Production are Associated with Progression from Barrett’s Esophagus to Esophageal Adenocarcinoma. bioRxiv. 2023. doi:10.1101/2023.06.27.546733

16. Kusters JG, van Vliet AHM, Kuipers EJ. Pathogenesis of Helicobacter pylori infection. Clin Microbiol Rev. 2006;19: 449–490.

17. Islami F, Kamangar F. Helicobacter pylori and esophageal cancer risk: a meta-analysis. Cancer Prev Res (Phila). 2008;1: 329–338.

18. Erőss B, Farkas N, Vincze Á, Tinusz B, Szapáry L, Garami A, et al. Helicobacter pylori infection reduces the risk of Barrett’s esophagus: A meta-analysis and systematic review. Helicobacter. 2018;23: e12504.

19. Du Y-L, Duan R-Q, Duan L-P. Helicobacter pylori infection is associated with reduced risk of Barrett’s esophagus: a meta-analysis and systematic review. BMC Gastroenterol. 2021;21: 459.

20. Kong CY, Kroep S, Curtius K, Hazelton WD, Jeon J, Meza R, et al. Exploring the recent trend in esophageal adenocarcinoma incidence and mortality using comparative simulation modeling. Cancer Epidemiol Biomarkers Prev. 2014;23: 997–1006.

21. Gall A, Fero J, McCoy C, Claywell BC, Sanchez CA, Blount PL, et al. Bacterial Composition of the Human Upper Gastrointestinal Tract Microbiome Is Dynamic and Associated with Genomic Instability in a Barrett’s Esophagus Cohort. PLoS One. 2015;10: e0129055.

22. Wiklund A-K, Santoni G, Yan J, Radkiewicz C, Xie S, Birgisson H, et al. Risk of Esophageal Adenocarcinoma After Helicobacter pylori Eradication Treatment in a Population-Based Multinational Cohort Study. Gastroenterology. 2024;167: 485–492.e3.

23. Sieber M, Pita L, Weiland-Bräuer N, Dirksen P, Wang J, Mortzfeld B, et al. Neutrality in the metaorganism. PLoS Biol. 2019;17: e3000298.

24. Venkataraman A, Bassis CM, Beck JM, Young VB, Curtis JL, Huffnagle GB, et al. Application of a neutral community model to assess structuring of the human lung microbiome. MBio. 2015;6. doi:10.1128/mBio.02284-14

25. Leung MHY, Tong X, Bastien P, Guinot F, Tenenhaus A, Appenzeller BMR, et al. Changes of the human skin microbiota upon chronic exposure to polycyclic aromatic hydrocarbon pollutants. Microbiome. 2020;8: 100.

26. Kim H-J, Kim H, Kim JJ, Myeong NR, Kim T, Park T, et al. Fragile skin microbiomes in megacities are assembled by a predominantly niche-based process. Sci Adv. 2018;4: e1701581.

27. Peter S, Pendergraft A, VanDerPol W, Wilcox CM, Kyanam Kabir Baig KR, Morrow C, et al. Mucosa-Associated Microbiota in Barrett’s Esophagus, Dysplasia, and Esophageal Adenocarcinoma Differ Similarly Compared With Healthy Controls. Clin Transl Gastroenterol. 2020;11: e00199.

28. Lopetuso LR, Severgnini M, Pecere S, Ponziani FR, Boskoski I, Larghi A, et al. Esophageal microbiome signature in patients with Barrett’s esophagus and esophageal adenocarcinoma. PLoS One. 2020;15: e0231789.

29. Zhou J, Shrestha P, Qiu Z, Harman DG, Teoh W-C, Al-Sohaily S, et al. Distinct Microbiota Dysbiosis in Patients with Non-Erosive Reflux Disease and Esophageal Adenocarcinoma. J Clin Med. 2020;9. doi:10.3390/jcm9072162

30. Liu N, Ando T, Ishiguro K, Maeda O, Watanabe O, Funasaka K, et al. Characterization of bacterial biota in the distal esophagus of Japanese patients with reflux esophagitis and Barrett’s esophagus. BMC Infect Dis. 2013;13: 130.

31. Pei Z, Bini EJ, Yang L, Zhou M, Francois F, Blaser MJ. Bacterial biota in the human distal esophagus. Proc Natl Acad Sci U S A. 2004;101: 4250–4255.

32. Paulson TG, Galipeau PC, Oman KM, Sanchez CA, Kuhner MK, Smith LP, et al. Somatic whole genome dynamics of precancer in Barrett’s esophagus reveals features associated with disease progression. Nat Commun. 2022;13: 2300.

33. Luebeck J, Ng AWT, Galipeau PC, Li X, Sanchez CA, Katz-Summercorn AC, et al. Extrachromosomal DNA in the cancerous transformation of Barrett’s oesophagus. Nature. 2023;616: 798–805.

34. ICGC/TCGA Pan-Cancer Analysis of Whole Genomes Consortium. Pan-cancer analysis of whole genomes. Nature. 2020;578: 82–93.

35. Ross-Innes CS, Becq J, Warren A, Cheetham RK, Northen H, O’Donovan M, et al. Whole-genome sequencing provides new insights into the clonal architecture of Barrett’s esophagus and esophageal adenocarcinoma. Nat Genet. 2015;47: 1038–1046.

36. Guccione C, Patel L, Tomofuji Y, McDonald D, Gonzalez A, Sepich-Poore G, et al. Incomplete human reference genomes can drive false sex biases and expose patient-identifying information in metagenomic data. Nat Commun. 2025 (in press, preprint available doi: 10.21203/rs.3.rs-4721159/v1).

37. Armstrong G, Martino C, Morris J, Khaleghi B, Kang J, DeReus J, et al. Swapping Metagenomics Preprocessing Pipeline Components Offers Speed and Sensitivity Increases. mSystems. 2022;7: e0137821.

38. Hillmann B, Al-Ghalith GA, Shields-Cutler RR, Zhu Q, Knight R, Knights D. SHOGUN: a modular, accurate and scalable framework for microbiome quantification. Bioinformatics. 2020;36: 4088–4090.

39. Zhu Q, Mai U, Pfeiffer W, Janssen S, Asnicar F, Sanders JG, et al. Phylogenomics of 10,575 genomes reveals evolutionary proximity between domains Bacteria and Archaea. Nat Commun. 2019;10: 5477.

40. Sepich-Poore GD, McDonald D, Kopylova E, Guccione C, Zhu Q, Austin G, et al. Robustness of cancer microbiome signals over a broad range of methodological variation. Oncogene. 2024;43: 1127–1148.

41. Steinegger M, Salzberg SL. Terminating contamination: large-scale search identifies more than 2,000,000 contaminated entries in GenBank. Genome Biol. 2020;21: 115.

42. Schneider VA, Graves-Lindsay T, Howe K, Bouk N, Chen H-C, Kitts PA, et al. Evaluation of GRCh38 and de novo haploid genome assemblies demonstrates the enduring quality of the reference assembly. Genome Res. 2017;27: 849–864.

43. Nurk S, Koren S, Rhie A, Rautiainen M, Bzikadze AV, Mikheenko A, et al. The complete sequence of a human genome. Science. 2022;376: 44–53.

44. Liao W-W, Asri M, Ebler J, Doerr D, Haukness M, Hickey G, et al. A draft human pangenome reference. Nature. 2023;617: 312–324.

45. Human Microbiome Project Consortium. Structure, function and diversity of the healthy human microbiome. Nature. 2012;486: 207–214.

46. Human Microbiome Project Consortium. A framework for human microbiome research. Nature. 2012;486: 215–221.

47. Hakim D, Wandro S, Zengler K, Zaramela LS, Nowinski B, Swafford A, et al. Zebra: Static and Dynamic Genome Cover Thresholds with Overlapping References. mSystems. 2022;7: e0075822.

48. Blanco-Míguez A, Beghini F, Cumbo F, McIver LJ, Thompson KN, Zolfo M, et al. Extending and improving metagenomic taxonomic profiling with uncharacterized species using MetaPhlAn 4. Nat Biotechnol. 2023;41: 1633–1644.

49. Sczyrba A, Hofmann P, Belmann P, Koslicki D, Janssen S, Dröge J, et al. Critical Assessment of Metagenome Interpretation-a benchmark of metagenomics software. Nat Methods. 2017;14: 1063–1071.

50. Meyer F, Fritz A, Deng Z-L, Koslicki D, Lesker TR, Gurevich A, et al. Critical Assessment of Metagenome Interpretation: the second round of challenges. Nat Methods. 2022;19: 429–440.

51. Bolyen E, Rideout JR, Dillon MR, Bokulich NA, Abnet CC, Al-Ghalith GA, et al. Reproducible, interactive, scalable and extensible microbiome data science using QIIME 2. Nat Biotechnol. 2019;37: 852–857.

52. Martino C, Morton JT, Marotz CA, Thompson LR, Tripathi A, Knight R, et al. A Novel Sparse Compositional Technique Reveals Microbial Perturbations. mSystems. 2019;4. doi:10.1128/mSystems.00016-19

53. Zapién-Campos R, Sieber M, Traulsen A. The effect of microbial selection on the occurrence-abundance patterns of microbiomes. J R Soc Interface. 2022;19: 20210717.

54. Hubbell SP. The Unified Neutral Theory of Biodiversity and Biogeography (MPB-32). Princeton University Press; 2001.

55. Yamamoto H, Bai YQ, Yuasa Y. Homeodomain protein CDX2 regulates goblet-specific MUC2 gene expression. Biochem Biophys Res Commun. 2003;300: 813–818.

56. Henihan RD, Stuart RC, Nolan N, Gorey TF, Hennessy TP, O’Morain CA. Barrett’s esophagus and the presence of Helicobacter pylori. Am J Gastroenterol. 1998;93: 542–546.

57. Xie F-J, Zhang Y-P, Zheng Q-Q, Jin H-C, Wang F-L, Chen M, et al. Helicobacter pylori infection and esophageal cancer risk: an updated meta-analysis. World J Gastroenterol. 2013;19: 6098–6107.

58. Mathieu LN, Kanarek NF, Tsai H-L, Rudin CM, Brock MV. Age and sex differences in the incidence of esophageal adenocarcinoma: results from the Surveillance, Epidemiology, and End Results (SEER) Registry (1973-2008). Dis Esophagus. 2014;27: 757–763.

59. Baker JL, Mark Welch JL, Kauffman KM, McLean JS, He X. The oral microbiome: diversity, biogeography and human health. Nat Rev Microbiol. 2024;22: 89–104.

60. Paczosa MK, Mecsas J. Klebsiella pneumoniae: Going on the Offense with a Strong Defense. Microbiol Mol Biol Rev. 2016;80.

61. Yadukumar L, Aslam H, Ahmed K, Iskander P, Sajid K, Syed O, et al. A Rare Case of Acute Esophageal Necrosis Precipitated by Klebsiella Pneumoniae. Gastro Hep Adv. 2023;2: 827–829.

62. Ren J-M, Zhang X-Y, Liu S-Y. Neisseria mucosa - A rare cause of peritoneal dialysis-related peritonitis: A case report. World J Clin Cases. 2023;11: 3311–3316.

63. Abdelsalam NA, Hegazy SM, Aziz RK. The curious case of Prevotella copri. Gut Microbes. 2023;15: 2249152.

64. Yu C, Zhou B, Xia X, Chen S, Deng Y, Wang Y, et al. Prevotella copri is associated with carboplatin-induced gut toxicity. Cell Death Dis. 2019;10: 714.

65. Scher JU, Sczesnak A, Longman RS, Segata N, Ubeda C, Bielski C, et al. Expansion of intestinal Prevotella copri correlates with enhanced susceptibility to arthritis. Elife. 2013;2: e01202.

66. Liu X, Li: LXS, Ji F, Mei Y, Cheng Y, Liu F, et al. Alterations of gastric mucosal microbiota across different stomach microhabitats in a cohort of 276 patients with gastric cancer. EBioMedicine. 2019;40: 336–348.

67. Gunathilake MN, Lee J, Choi IJ, Kim Y-I, Ahn Y, Park C, et al. Association between the relative abundance of gastric microbiota and the risk of gastric cancer: a case-control study. Sci Rep. 2019;9: 13589.

