## Supplementary Information for "Community assembly modeling of microbial evolution within Barrett’s esophagus and esophageal adenocarcinoma"

### Supplementary Methods

#### *micov coverage filter methods*

To identify any unreliable taxa mapping in the main results, we employed an aggregate MIcrobiome COVerage tool (micov). Micov computes coverage dispersions across many microbial genomes and samples to filter out taxa with low genomic coverage from further analysis. Micov takes SAM/BAM or BED3 files across multiple samples as input, which contain reads aligned to a database of reference genomes (as is the case when using a microbial database like Web of Life). Using the CIGAR string, and subject start position from the alignment data, it calculates per-sample genome coverage (including per-position), as well as metrics determining if the reads mapped to the taxa are represented across the entire length of the taxa or just a small region. Taxa that do not have sufficient coverage from their mapped reads across the length of their genome, as determined based on criteria below, are dropped from the taxonomic table. We note that genome, in this case, refers to consecutive regions of fragmented assemblies that may not reflect the true genome structure.

Specifically, for each *genome_id,* user input to micov includes *genome_length* (the length of the genome) and micov then calculates: *base_pair_covered* (total number of base pairs in the genome for which any read aligns to), *percent_covered* (*base_pair_covered* / *genome_length*), and a data frame of all aligned reads to the *genome_id* with the following columns: *sample_id*, *start_position* and *stop_position* (for read location in the *genome_id*). Then micov creates bins of length 10,000 base pairs across the *genome_length*, for example: bin 1 : (0, 10,000), bin 2 : (10,001, 20,000*)*, … bin n : (*genome_length* - 10,000, *genome_length*). It then creates a *grouped_bins* data frame that starts with all bins that contain at least a single read and joins together any adjacent bins. The resulting data frame contains the following columns: *start_position* and *stop_position* of the entire grouped bin, *reads_list* (a list of unique IDs for each read/hit within this bin), and *samples_list* (a list of all samples that have at least one read in this bin). These *grouped_bins* represent genome regions for which reads from our samples are aligned. For confident taxon identification in the dataset, we would expect a few very large *grouped_bins* in the case of high sequencing depth for that taxon or multiple *grouped_bins* distributed broadly across the length of the genome in the case of low sequencing depth for that taxon.

Before further filtering, micov identifies whether each *genome_id* (taxon) has low or high sequencing depth. If the combined coverage of the *genome_id* across all samples (*percent_covered*) is greater than 10%, the *genome_id* is considered high coverage and automatically **passes**. The 10% threshold cutoff is based on Zebra [1]. Thus the largest possible bin following this threshold would still only cover <10% of the genome. For remaining low-coverage genomes (<10% total coverage), we use the following thresholds to determine if the genomic coverage on a taxon is sufficient in this order:

1. If there are fewer than five counts as determined by woltka for the *genome_id* in the taxonomy table, we do not have enough information to be confident this is a real microbial count, and therefore the *genome_id* **fails**.

2. If there are less than 30 total reads for the *genome_id*, similar to the argument in (1), we do not have enough information to determine if this is a true count, and therefore the *genome_id* **fails**. We note that reads and counts in the taxonomy table are not always equal due to the multi-mapping nature of the SHOGUN [2] pipeline, with up to 16 taxonomic mappings for each read.

3. If there are fewer than four bins in the *grouped_bins*, we do not have enough diversity in locations covered across the genome to be confident we have identified a correct taxa, and thus the *genome_id* **fails**. Note, as explained above, this is following the removal of high coverage taxa which may have one very large grouped bin. Listed as criteria A in main text.

4. We consider a subset of samples that only each contribute reads to a single *grouped_bin* for the particular *genome_id* (thus called *single_bin_samples*). Then, looking at each aligned read to the *genome_id*, we determine if the read is from a sample in the *single_bin_samples* list. If greater than 75% of reads fall into this category, we are not confident that any sample truly identifies this taxon since all samples are mapping to such a small proportion of the genome, and thus, the *genome_id* **fails**. Note that in the case of BE non-cancer outcomes (NCO) and cancer outcomes (CO), we have low sequencing depth due to epithelial isolation preparation, so we did not use this threshold. Listed as criteria B in the main text.

5. Using the *grouped_bins* data frame, we identify the top three bins with the largest number of samples in their *samples_list*. We then calculate the total number of reads in these bins with their *reads_lists*. If greater than 75% of the total reads fall into these top 3 bins, then the majority of hits are not evenly spread across the genome, and therefore, we are not confident we have identified the correct taxa, so this *genome_id* **fails**. As with (4), in the case of BE NCO and CO, we have low sequencing depth due to epithelial isolation preparation, so we did not use this threshold. Listed as criteria C in main text.

All *genome_ids* that have >10% coverage or satisfy criteria 1-5 above (or 1-3 in the case of BE NCO/CO cohorts) are considered sufficiently mapped and are used for downstream analyses. All *genome_ids* that have <10% coverage and fail any of the 1-5 criteria are dropped from the taxonomy table and not used in our analysis.

### Supplementary Tables

**Supplementary Table S1. Description of patient datasets.** Number of samples pre and post-rarefaction, rarefaction level of whole genome sequencing (WGS) data, sampling technique, and citation provided for each dataset provided that have additional details on WGS processing. Human reference genomes GRCh38.p14 and CHM13-T2Tv2.0 were used for host depletion in all WGS datasets (see Materials and methods in main text).

| Disease type | Sample type | # of samples | # of samples following rarefaction | Rarefaction depth | Sampling technique | Ref |
|---|---|---|---|---|---|---|
| Healthy esophagus | Esophageal brushing | 50 | 42 | 80,000 | During endoscopy, an esophageal brushing was taken and extreme care was taken to ensure the (non-reusable) brush was not contaminated by saliva. | [3] |
| GERD esophagus | | 29 | 26 | 80,000 | | |
| BE Non-Cancer Outcome | Isolated epithelium from esophageal tissue biopsy | 80 | 63 | 14,000 | During endoscopy, endoscopic biopsies were taken from the BE segment and stored fresh-frozen. | [4] |
| BE Cancer Outcome | | 80 | 59 | 14,000 | | |
| EAC: Cohort 1 | Esophageal tumor tissue | 83 | 68 | 150,000 | Tissue underwent cryopreservation in liquid nitrogen (dead tissue) and kept frozen at -70°C. | [5] |
| EAC: Cohort 2 | Esophageal tumor tissue | 23 | 19 | 150,000 | Tissue samples were obtained from surgical resection, endoscopic ultrasound or endoscopic mucosal resection. They were snap-frozen in liquid nitrogen immediately after collection and stored at -80°C. | [6] |

**Supplementary Table S2. Top 10 non-neutral microbial species in non-progressor (NCO; non-cancer outcome) BE samples.** Difference values are the residuals of data for the specific microbial species to the expected occurrence frequency obtained from the best-fit neutral prediction. Species in bold text were found in the top 10 lists for both NCO and cancer outcome patients. Species colored in red were included in 5 human-associated microbial surveys [7].

| Difference from Neutral Model in NCO BE | Microbial Species | Relevant Research | Citation |
|---|---|---|---|
| 0.874348995 | *Klebsiella pneumoniae* | *K. pneumoniae* is commonly found in the human gut but can cause severe infection across the human body with high mortality rates, especially in immunocompromised patients. It has been identified as a primary pathogen for pyogenic liver abscesses (PLA), and patients with *K. pneumoniae* PLA had higher rates of colorectal cancer (CRC) which may be due to the polyketide synthases (PKS) genes within both *K. pneumoniae* and *E. coli*. A case study also linked *K. pneumoniae* with Acute Esophageal Necrosis. | [8–12] |
| 0.815623023 | *Helicobacter pylori* | *H. pylori* is well known to cause chronic inflammation, gastric ulcer disease, and gastric cancer, especially if the strain expresses CagA. Additionally, *H. pylori* infection has been associated with a decreased risk of both esophageal squamous cell carcinoma (ESCC) and esophageal adenocarcinoma (EAC). | [13–15] |
| 0.645910100 | ***Shigella dysenteriae*** | Closely related to *E. coli*, *S. dysenteriae* is a normal microflora of the human gut, but can also cause serious illness when it invades and damages epithelial cells in the gastrointestinal tract leading to bloody diarrhea and hemolytic uremic syndrome. Depending on the strain, *S. dysenteriae* also contains Shiga toxin (Stx), an extremely potent bacterial toxin. | [16,17] |
| 0.570354669 | ***Shigella flexneri*** | Similar to *S. dysenteriae, S. flexneri* leads to diarrhea mixed with blood and mucus, especially in children under five or those with malnutrition. In rare cases, it can lead to neurological complications such as encephalopathy. One study demonstrated that *S. flexneri* may even have an anti-proliferative effect on pancreatic cancer cells through apoptosis by upregulating the pro-apoptotic gene Bax and downregulating the anti-apoptotic gene bcl-2. | [18–20] |
| 0.410072979 | *Propionibacterium namnetense* | *P. namnetense* (now known as *Cuitbacterium namnetense*) is a relatively new bacteria that was isolated from a tibia former external fixator. It has been linked with a few human infections in rare cases, | [21–24] |

| | | | |
|---|---|---|---|
| | | including osteosynthetic cervical spine infection and liver abscesses. | |
| 0.396825294 | *Porphyromonas somerae* | *P. somerae* was originally isolated from patients with chronic skin and soft tissue infections. It has been found in patients with endometrial cancer and may potentially have a role in its progression and serve as a biomarker of the disease. | [25–30] |
| 0.380531138 | *Propionibacterium humerusii* | *P. humerusii* is a minimally studied bacteria that is often confused with *P. acnes* (*Cutibacterium acnes*). | [31] |
| 0.338339349 | *Neisseria cinerea* | *N. cinerea* is often found in the human oropharynx and is very rarely associated with serious infection in humans. It is extremely similar to both *N. gonorrhoeae* and *N. mucosa* which are more well studied in humans. | [32,33] |
| 0.333333333 | ***Neisseria mucosa*** | *N. mucosa* is found in oral and nasal mucosa as well as the upper respiratory tract of humans. It can occasionally cause upper respiratory issues or diseases such as endocarditis in immunocompromised patients. | [34] |
| 0.298624304 | *Corynebacterium kroppenstedtii* | *C. kroppenstedtii* is a microbe that has been mainly isolated from female patients specifically in breast abscesses. This microbe has been linked to both granulomatous mastitis and potentially psychiatric illnesses. Additionally, it can cause bloodstream infections in rare cases. | [35–37] |

**Supplementary Table S3. Top 10 non-neutral microbial species in progressor (CO; cancer outcome) BE samples.** Values are the residuals of data for the specific microbial species to the expected occurrence frequency obtained from the best-fit neutral prediction. Species in bold text were found in the top 10 lists for both non-cancer outcome and CO patients. Species colored in red were included in 5 human-associated microbial surveys [7].

| Difference from Neutral Model in CO BE | Microbial Species | Relevant Research | Citation |
|---|---|---|---|
| 0.755032266 | ***Shigella dysenteriae*** | Closely related to *E. coli*, *S. dysenteriae* is a normal microflora of the human gut, but can also cause serious illness when it invades and damages epithelial cells in the gastrointestinal tract leading to bloody diarrhea and hemolytic uremic syndrome. Depending on the strain, *S. dysenteriae* also contains Shiga toxin (Stx), an extremely potent bacterial toxin. | [16,17] |
| 0.750225422 | *Lactobacillus fermentum* | A very common non-pathogenic, human-origin microbe that is used commonly in food fermentation and has been shown to have adaptive immune processes in diverse inflammatory diseases. Depending on the strain, it has been linked to tumor metastasis inhibition in mice. | [38] |
| 0.688264102 | *Alishewanella agri* | *A. agri* is isolated from landfill soil and does not appear to have a human link. | [39] |
| 0.668818908 | *Pseudomonas mendocina* | *P. mendocina* has been isolated from water and soil and can rarely cause severe infection in humans. | [40] |
| 0.664854372 | ***Shigella flexneri*** | Similar to *S. dysenteriae, S. flexneri* leads to diarrhea mixed with blood and mucus, especially in children under five or those with malnutrition. In rare cases, it can lead to neurological complications such as encephalopathy. One study demonstrated that *S. flexneri* may even have an anti-proliferative effect on pancreatic cancer cells through apoptosis by upregulating the pro-apoptotic gene Bax and downregulating the anti-apoptotic gene bcl-2. | [18–20] |
| 0.498967543 | *Prevotella copri* | *P. copri* is an abundant member of the human gastrointestinal microbiome with potentially negative and positive effects on human health. Studies in mice found correlations with *P. copri* and worse intestinal mucositis and exacerbated colitis. | [41] |
| 0.456586353 | ***Neisseria mucosa*** | *N. mucosa* is found in the upper respiratory tract of humans and is occasionally linked with endocarditis and upper respiratory issues. | [34] |
| 0.411535177 | *Burkholderia ubonensis* | *B. ubonensis* is a bacterium that can cause opportunistic infection in patients with weak immune systems. | [42] |
| 0.407273134 | *Alishewanella sp. HH-ZS* | A recently discovered soil microbe that is not known to infect humans. | [43] |
| 0.403835897 | *Morococcus cerebrosus* | *M. cerebrosus* has been isolated from the human brain and supragingival plaque and calculus. | [44–46] |

### Supplementary Figures

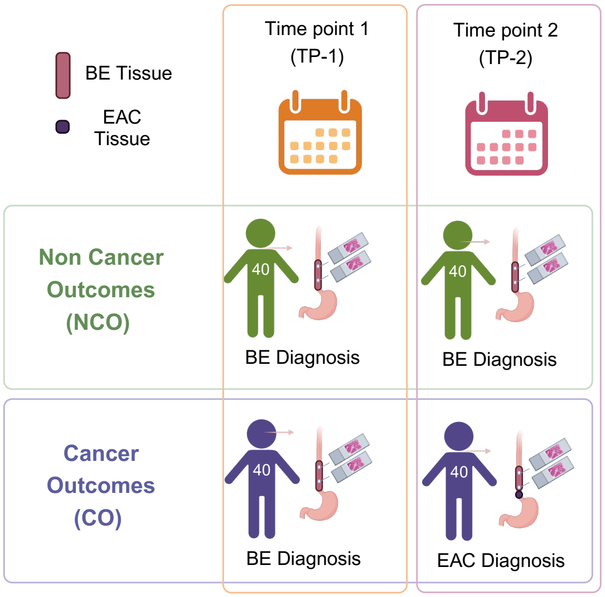


**Supplementary Figure S1. Barrett's esophagus (BE) case-control study design.** Paulson et al., study design for data collected from a total of 80 patients each with samples at two time points [4]. The patients were defined as Non-Cancer Outcomes (NCO; non-progressors) or Cancer Outcomes (CO; progressors to esophageal adenocarcinoma [EAC]). At each time point, an upper and lower tissue biopsy was collected from the patient's BE segment.

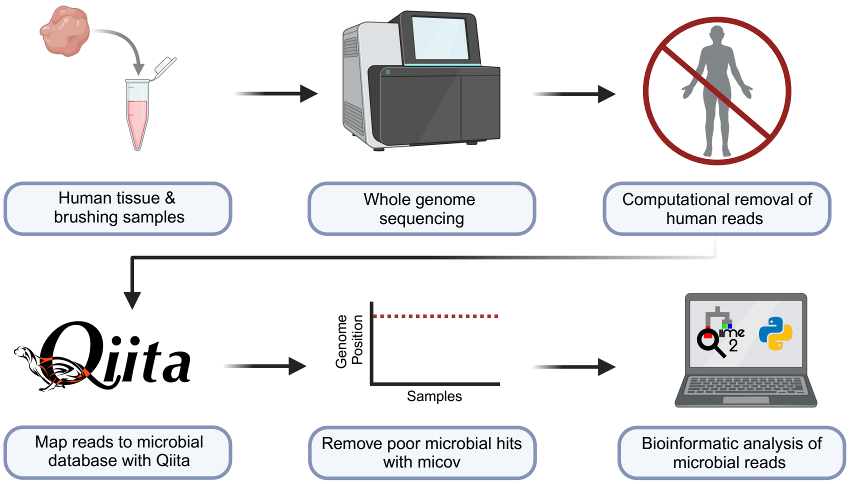


**Supplementary Figure S2. Pipeline of analysis from DNA extracted from human tissue samples to assignment of microbial DNA reads.** Human tissue samples were obtained by either biopsy or esophageal brushing, extracted DNA is prepared and whole genome sequenced, and then metagenomic data is bioinformatically depleted of human reads and mapped to microbial taxa. Poor microbial hits are removed and remaining microbial reads are used in downstream computational analyses.

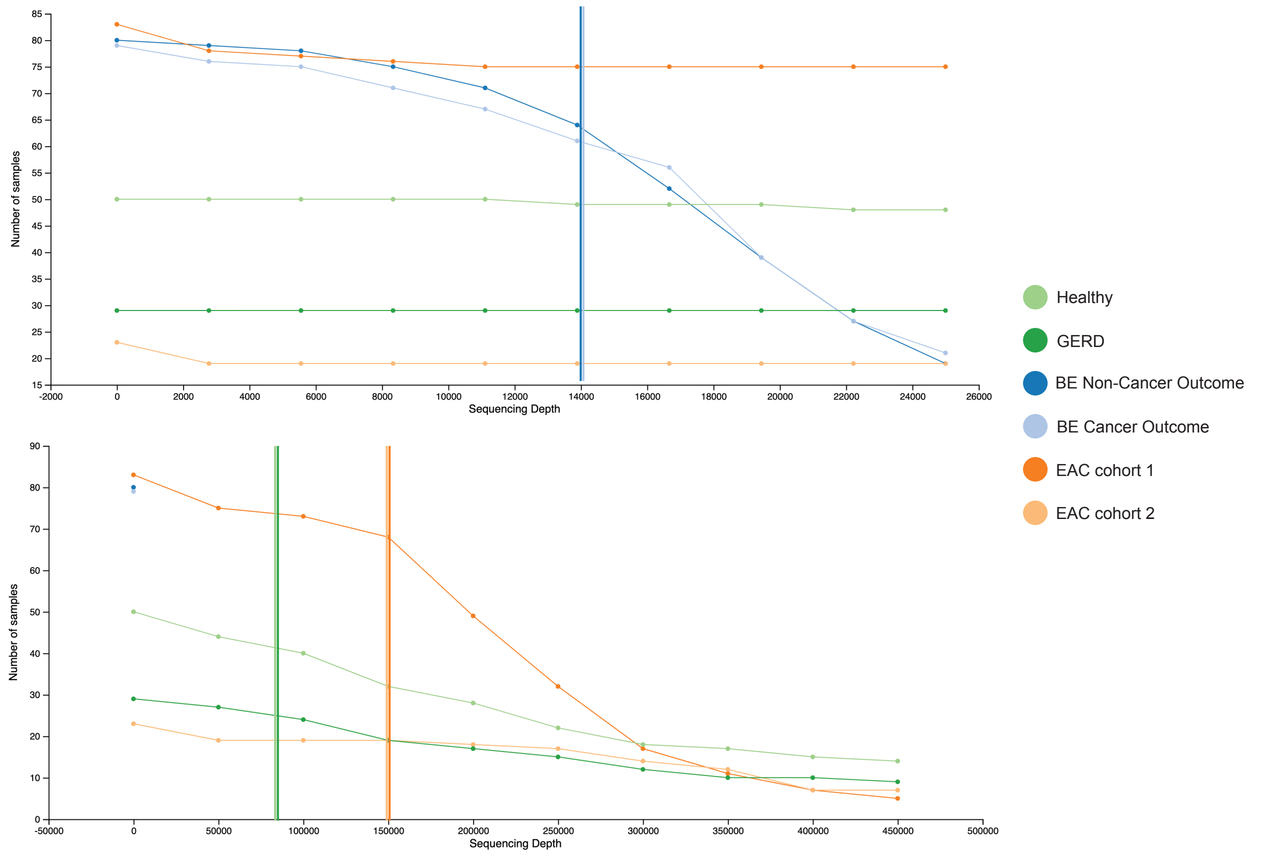


**Supplementary Figure S3. Alpha rarefaction curves for datasets analyzed in study.** Various sequencing depth subsets were calculated for each sample group analyzed. The colored vertical lines highlight the cutoff used in results for each independent dataset. The top graph shows sequencing depth from 0 to 24,000 read depth (capturing BE sample sequencing depths), and the bottom graph shows sequencing depth from 0 to 450,000 read depth.

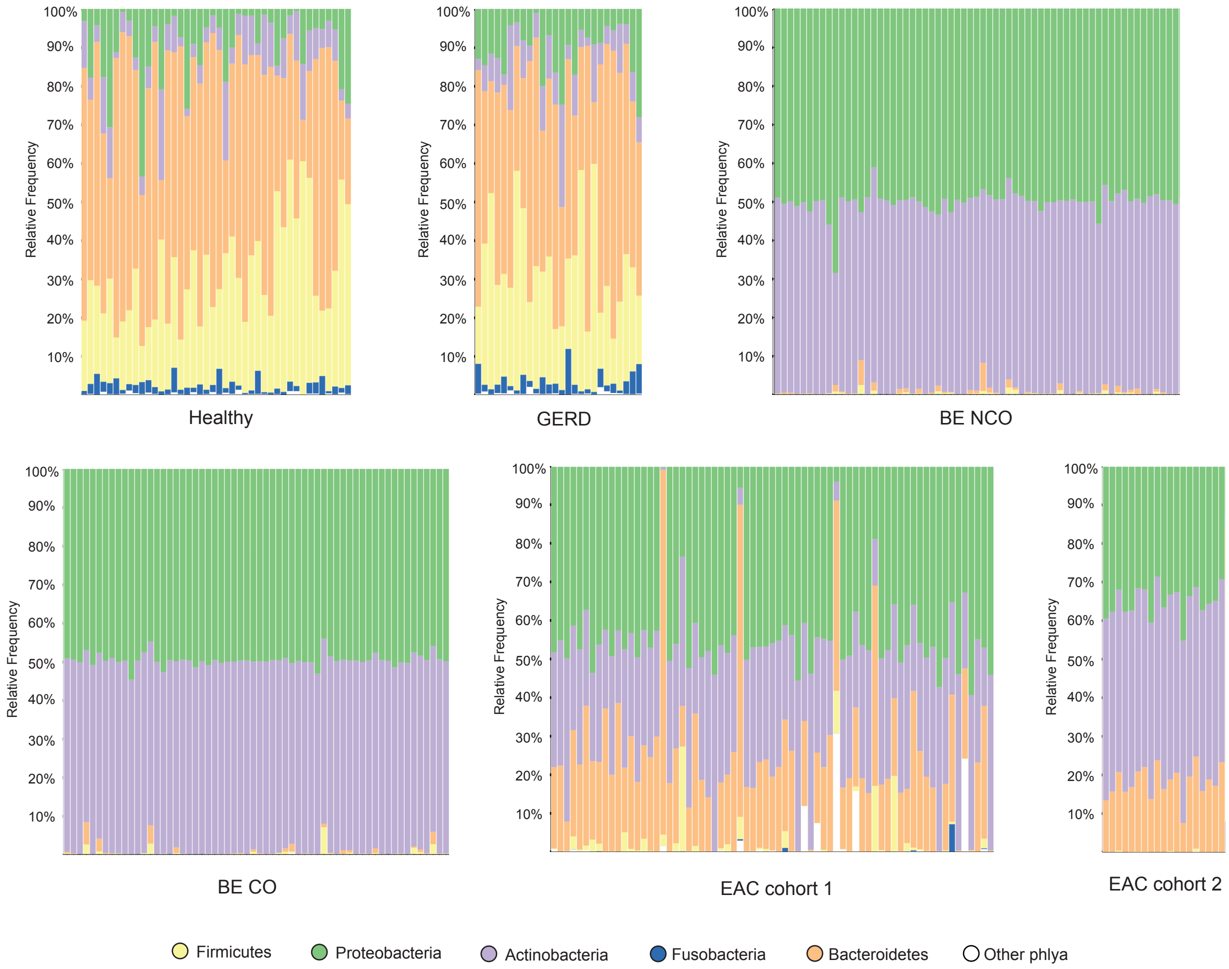


**Supplementary Figure S4. Most prevalent phylum level taxonomic abundances across disease stages.** Each vertical bar represents one sample. In the case of BE NCO and BE CO, only TP-1 data are shown (as included throughout main Results). Both samples (upper and lower) for each patient at TP-1 are displayed for the BE NCO and CO cohorts.

A

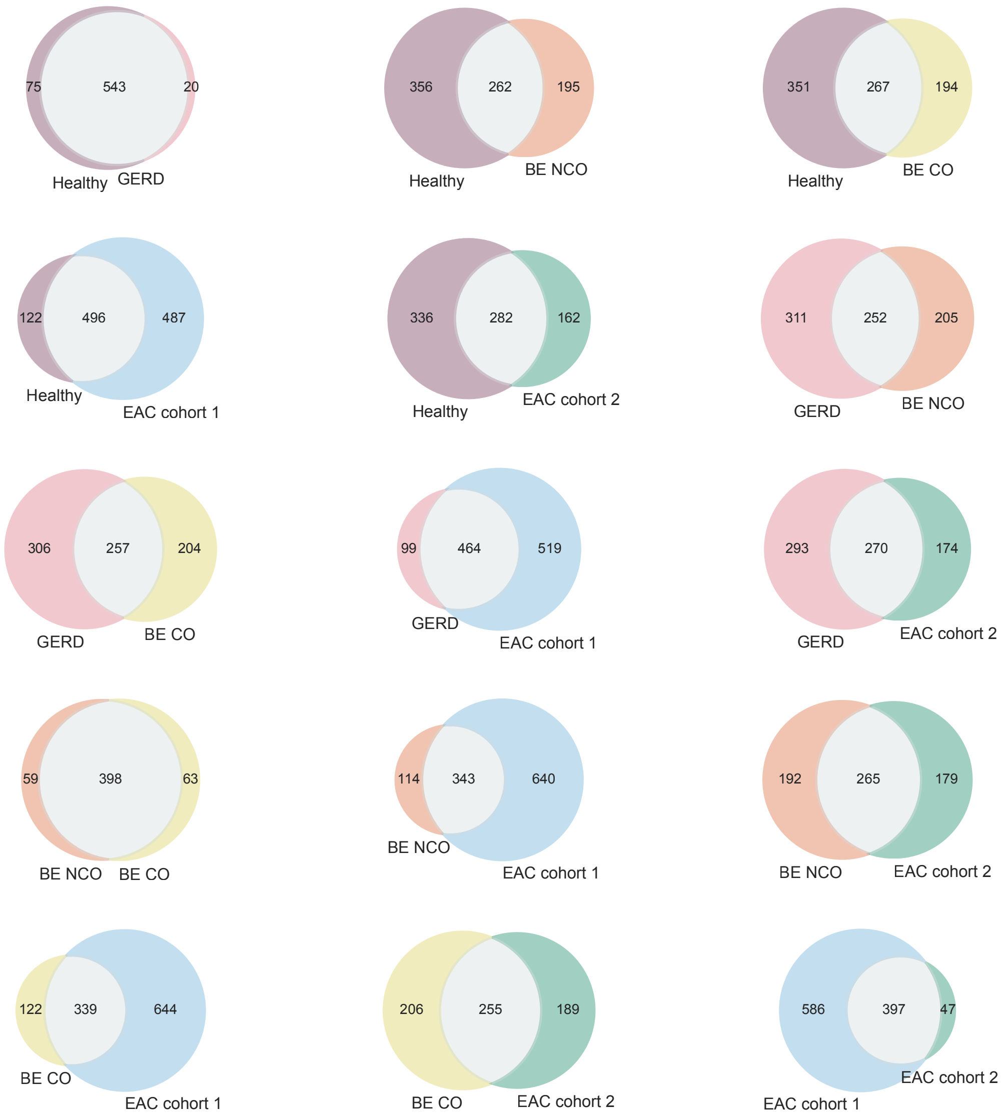


B

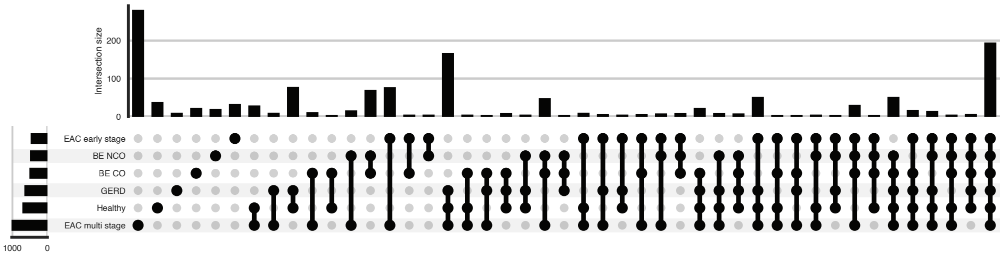


**Supplementary Figure S5. Comparison of microbial taxa identified across datasets. A.** Venn diagrams illustrate numbers of overlapping and unique microbial taxa quantified from rarefied data in cross-comparisons. **B**. UpSet plot of microbial taxa from rarefied datasets.

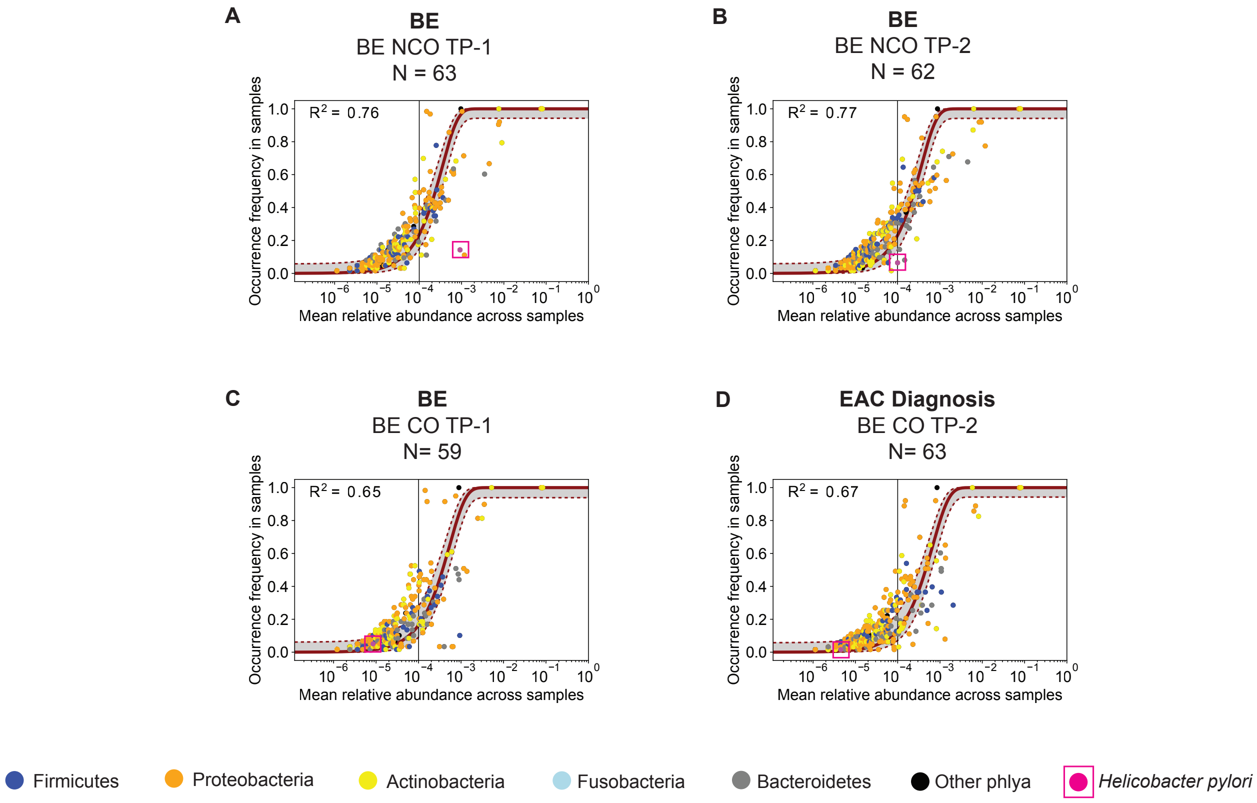


**Supplementary Figure S6. Community assembly dynamics of data from Paulson et al., at time point 2. A.** Same data as shown for BE NCO panel in Figure 3, with neutral model fit for BE NCO patients at TP-1. **B.** Same patients as in **A**, at time point 2 (TP-2) when patients still have BE. **C.** Same data as shown for BE CO panel in Figure 3, with neutral model fit for BE CO patients at TP-1. **D**. Same patients as in **C** at time point 2 (TP-2) but BE sample taken when patients have been diagnosed with EAC. The red curves represent the neutral model fit to data points representing unique taxa at the species level. The gray regions bound by red dotted lines represent the 95% bootstrap confidence intervals (obtained by resampling the hosts 100 times with replacement and refitting) and the $R^2$ value represents goodness of fit to the neutral model (see Materials and methods for details). The color of the data point represents the phyla of the microbial taxa indicated in the legend. *Helicobacter pylori* is highlighted in pink with a box outline. The rarefaction level used across all panels was 14,000.

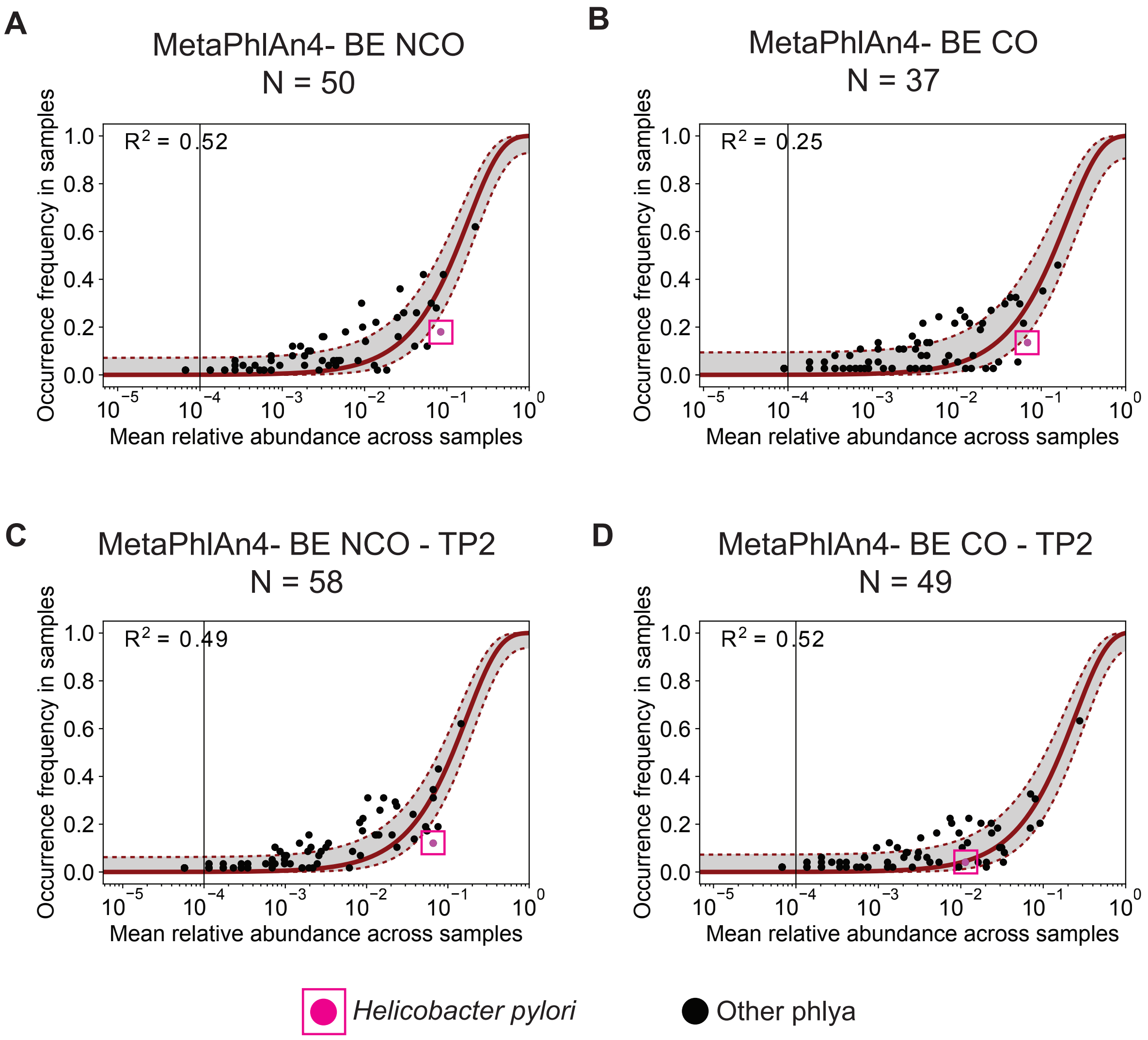


**Supplementary Figure S7. Community assembly dynamics of BE samples as analyzed with MetaPhlAn4.** Same patients were included as in Figure 3 (BE NCO) (**A**), Figure 3 (BE CO) (**B**), Supp. Figure S6B (**C**), Supp. Figure S6D (**D**), but using MetaPhlAn4 instead of the host depletion pipeline with Qiita and Qiime2 (Supplementary Figure S2). The gray regions bound by red dotted lines represent the 95% bootstrap confidence intervals (obtained by resampling the hosts 100 times with replacement and refitting) and the $R^2$ value represents goodness of fit to the neutral model (see Materials and methods for details). *Helicobacter pylori* is plotted in pink with a box outline.

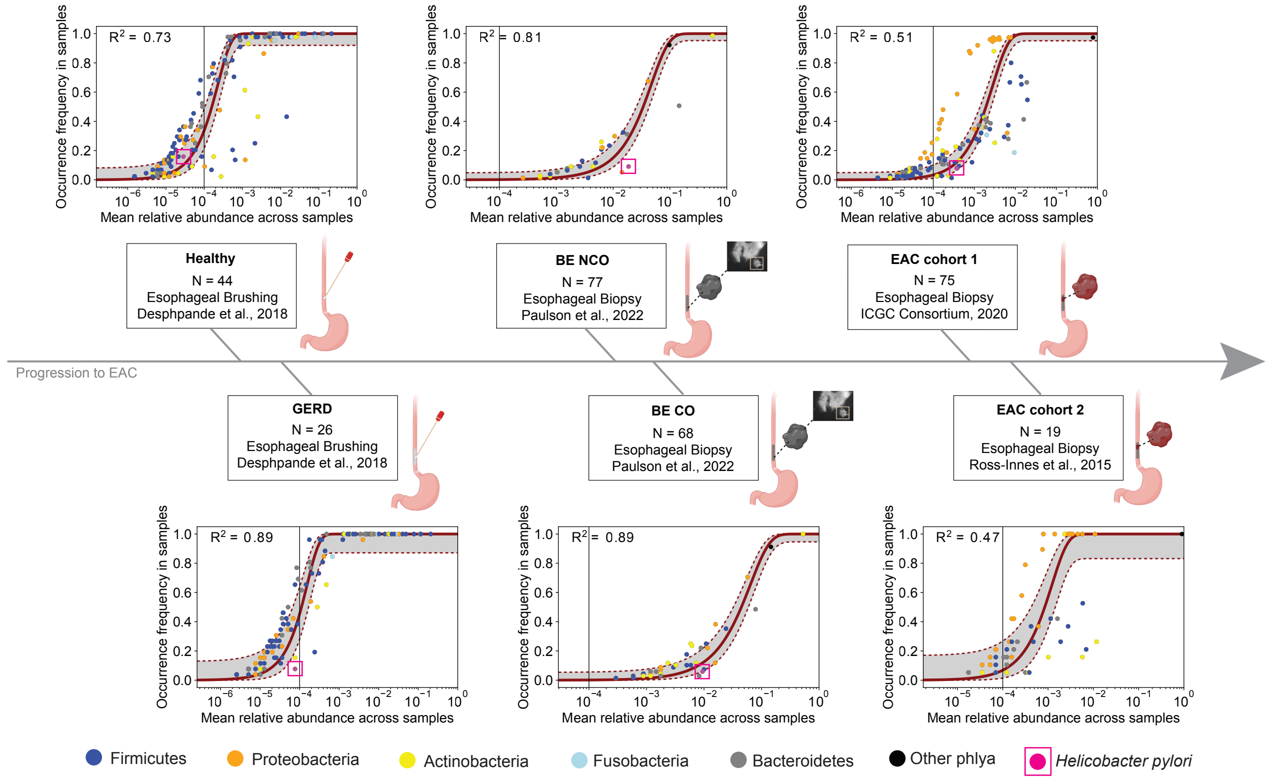


**Supplementary Figure S8. Community assembly dynamics in progression to EAC considering only taxa in a list of human-associated microbes.** This figure is similar to Figure 3, except identified microbes were subset to only microbes also found specifically in 5 human-associated microbial surveys [7] as well as the Web of Life - Clean database. As in Figure 3, plots are in order of disease state progression from healthy to EAC (see Supplementary Table S1 for patient/sample details). The red curves represent the neutral model fit to data points representing unique taxa at the species level. The gray regions bound by red dotted lines represent the 95% bootstrap confidence intervals (obtained by resampling the hosts 100 times with replacement and refitting) and the $R^2$ value represents goodness of fit to the neutral model (see Materials and methods for details). The color of the data point represents the phyla of the microbial taxa indicated in the legend. *Helicobacter pylori* is plotted in pink with a box outline. The rarefaction level used for each of the datasets were: Health/GERD: 15,000; BE NCO/CO 50; EAC cohorts: 3,000.

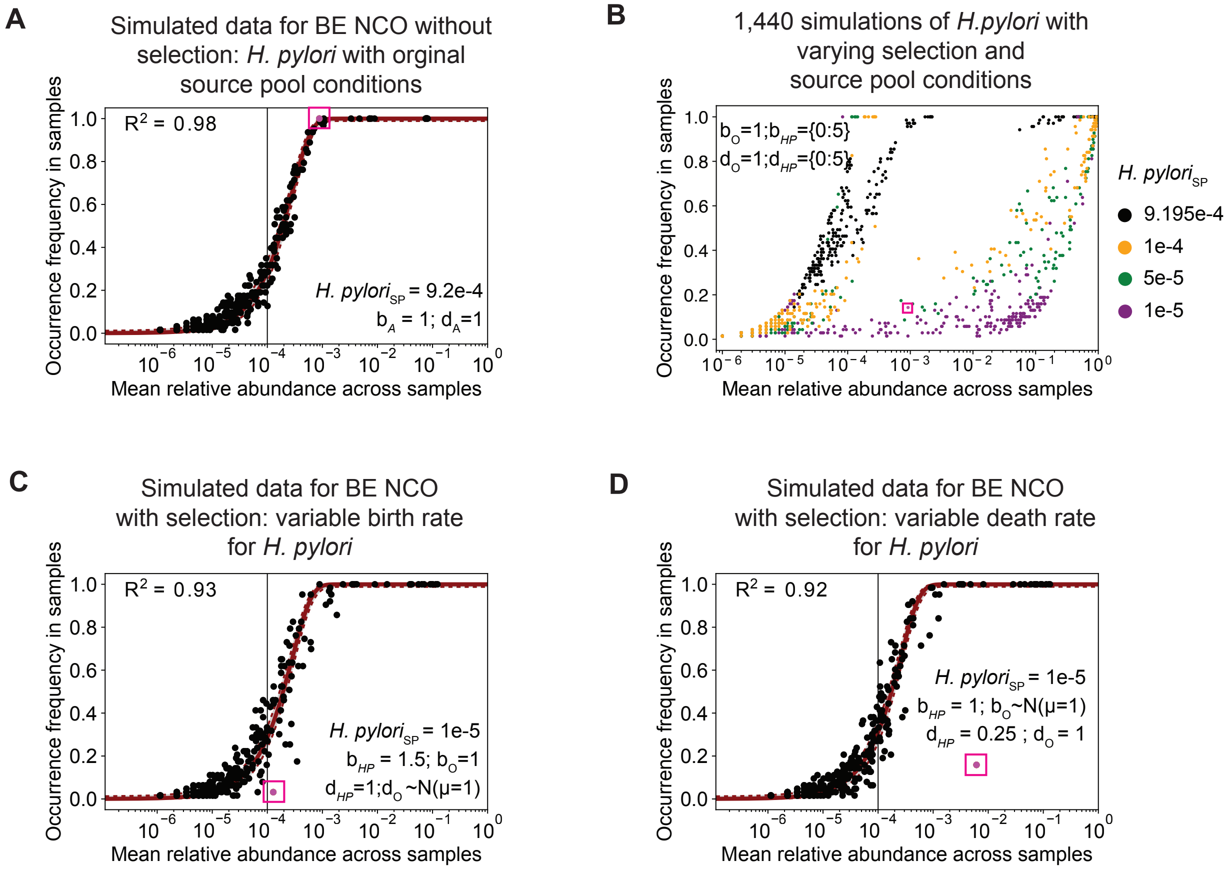


**Supplementary Figure S9. Additional simulation results for BE patients with non-cancer outcome (NCO). A.** Neutral simulation with data from the BE NCO patients using the source pool for *H. pylori* found in BE NCO. All ('A') microbes have an equal birth and death rate of 1. **B.** Varying occurrence-abundance locations of *H. pylori* after running the non-neutral simulation with a range of different birth rates (0-5) and death rates (0-5) as well as various source pool conditions for *H. pylori* (9.2e-4 [original position used in **A**], 1e-4, 5e-5, 1e-5). The pink box is the location of *H. pylori* in the BE NCO data. **C.** Non-neutral simulation for BE NCO data with *H. pylori* adjusted source pool prevalence (*H. pylori*$_{SP}$ =1e-5) and higher *H. pylori* birth rate than all other ('O') microbial taxa ($b_O$ =1). For the *H. pylori* taxon, the birth and death rates were set to 1.5 and 1, respectively. For all other microbes, the death rates were drawn independently from a normal distribution (mean=1, standard deviation=0.1). **D.** Non-neutral simulation for BE NCO data with *H. pylori* adjusted source pool prevalence (*H. pylori*$_{SP}$ =1e-5) and a lower death rate than all other microbial taxa ($d_O$ =1). For the *H. pylori* taxon, the birth and death rates were set to 1 and 0.25, respectively. For all other microbes, the birth rates were drawn independently from a normal distribution (mean=1, standard deviation=0.1). For **A, C-D**: The pink dot with outline indicates simulated data for *H. pylori*.

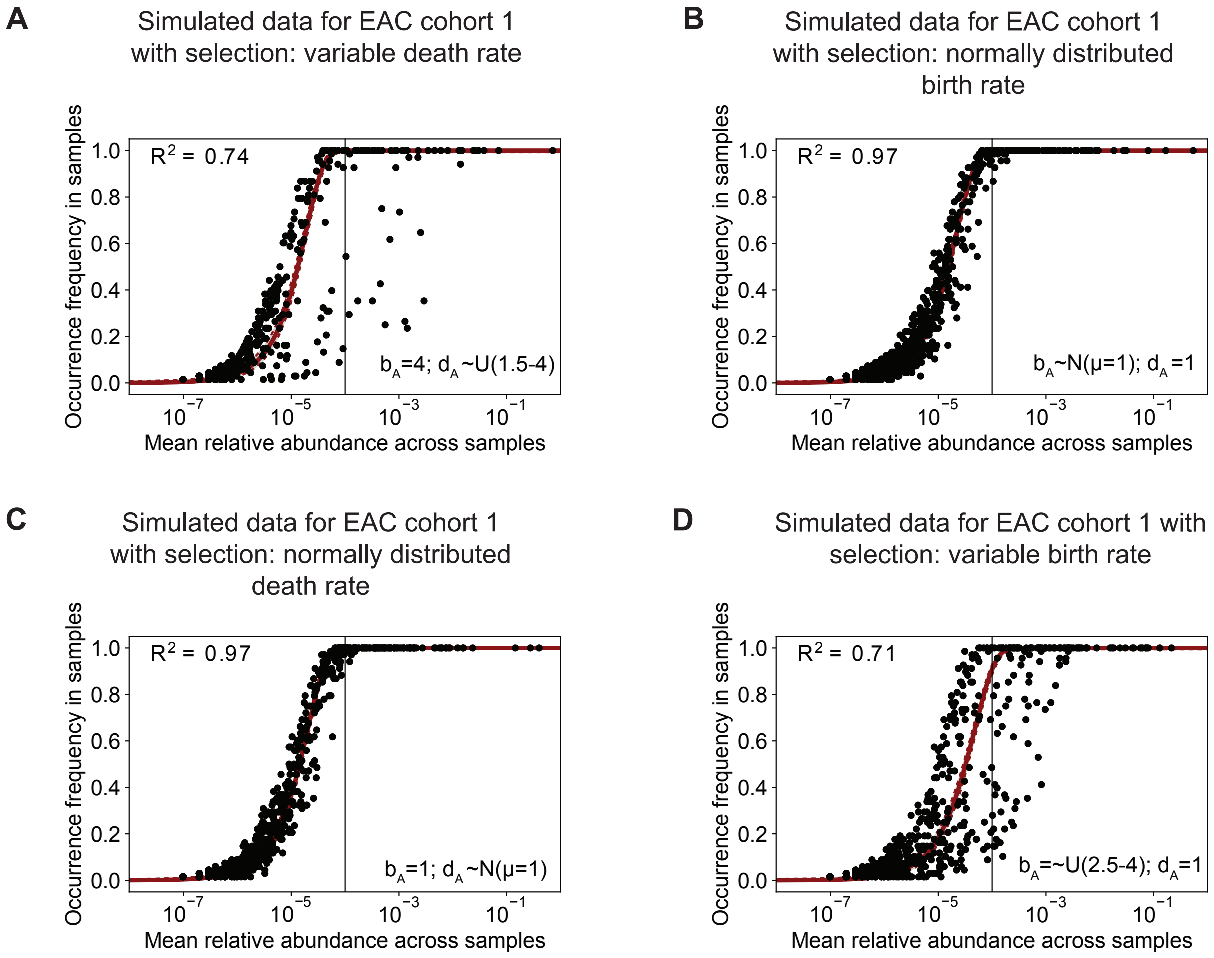


**Supplementary Figure S10. Additional simulation results for EAC cohort 1 patients. A.** Non-neutral model simulation with all microbes ('A') having variable death rates drawn independently from a uniform distribution ranging from 1.5 to 4, and a birth rate of 4. **B.** Non-neutral model simulation with all microbes having variable birth rates drawn independently from a normal distribution (mean=1, standard deviation= 0.1) and a death rate of 1. **C.** Non-neutral model simulation with all microbes having variable death rates drawn independently from a normal distribution (mean=1, standard deviation= 0.1) and a birth rate of 1. **D.** Non-neutral model simulation with all microbes having variable birth rates drawn independently from a uniform distribution ranging from 2.5 to 4 and a death rate of 1.

### Supplementary References


1. Hakim D, Wandro S, Zengler K, Zaramela LS, Nowinski B, Swafford A, et al. Zebra: Static and Dynamic Genome Cover Thresholds with Overlapping References. mSystems. 2022;7: e0075822.

2. Hillmann B, Al-Ghalith GA, Shields-Cutler RR, Zhu Q, Knight R, Knights D. SHOGUN: a modular, accurate and scalable framework for microbiome quantification. Bioinformatics. 2020;36: 4088–4090.

3. Deshpande NP, Riordan SM, Castaño-Rodríguez N, Wilkins MR, Kaakoush NO. Signatures within the esophageal microbiome are associated with host genetics, age, and disease. Microbiome. 2018;6: 227.

4. Paulson TG, Galipeau PC, Oman KM, Sanchez CA, Kuhner MK, Smith LP, et al. Somatic whole genome dynamics of precancer in Barrett's esophagus reveals features associated with disease progression. Nat Commun. 2022;13: 2300.

5. ICGC/TCGA Pan-Cancer Analysis of Whole Genomes Consortium. Pan-cancer analysis of whole genomes. Nature. 2020;578: 82–93.

6. Ross-Innes CS, Becq J, Warren A, Cheetham RK, Northen H, O'Donovan M, et al. Whole-genome sequencing provides new insights into the clonal architecture of Barrett's esophagus and esophageal adenocarcinoma. Nat Genet. 2015;47: 1038–1046.

7. Battaglia TW, Mimpen IL, Traets JJH, van Hoeck A, Zeverijn LJ, Geurts BS, et al. A pan-cancer analysis of the microbiome in metastatic cancer. Cell. 2024;187: 2324-2335.e19.

8. Strakova N, Korena K, Karpiskova R. Klebsiella pneumoniae producing bacterial toxin colibactin as a risk of colorectal cancer development - A systematic review. Toxicon. 2021;197: 126–135.

9. Lederman ER, Crum NF. Pyogenic liver abscess with a focus on Klebsiella pneumoniae as a primary pathogen: an emerging disease with unique clinical characteristics. Am J Gastroenterol. 2005;100: 322–331.

10. Huang W-K, Chang JW-C, See L-C, Tu H-T, Chen J-S, Liaw C-C, et al. Higher rate of colorectal cancer among patients with pyogenic liver abscess with Klebsiella pneumoniae than those without: an 11-year follow-up study. Colorectal Dis. 2012;14: e794-801.

11. Kaur CP, Iyadorai T, Sears C, Roslani AC, Vadivelu J, Samudi C. Presence of Polyketide Synthase (PKS) Gene and Counterpart Virulence Determinants in Klebsiella pneumoniae Strains Enhances Colorectal Cancer Progression In-Vitro. Microorganisms. 2023;11. doi:10.3390/microorganisms11020443

12. Yadukumar L, Aslam H, Ahmed K, Iskander P, Sajid K, Syed O, et al. A Rare Case of Acute Esophageal Necrosis Precipitated by Klebsiella Pneumoniae. Gastro Hep Adv. 2023;2: 827–829.

13. Wroblewski LE, Peek RM Jr, Wilson KT. Helicobacter pylori and gastric cancer: factors that modulate disease risk. Clin Microbiol Rev. 2010;23: 713–739.

14. Baj J, Forma A, Sitarz M, Portincasa P, Garruti G, Krasowska D, et al. Virulence Factors-Mechanisms of Bacterial Pathogenicity in the Gastric Microenvironment. Cells. 2020;10. doi:10.3390/cells10010027

15. Xie F-J, Zhang Y-P, Zheng Q-Q, Jin H-C, Wang F-L, Chen M, et al. Helicobacter pylori infection and esophageal cancer risk: an updated meta-analysis. World J Gastroenterol. 2013;19: 6098–6107.

16. Pakbin B, Brück WM, Brück TB. Molecular Mechanisms of Shigella Pathogenesis; Recent Advances. Int J Mol Sci. 2023;24. doi:10.3390/ijms24032448

17. Melton-Celsa AR. Shiga Toxin (Stx) Classification, Structure, and Function. Microbiol Spectr. 2014;2: EHEC-0024-2013.

18. Nisa I, Qasim M, Yasin N, Ullah R, Ali A. Shigella flexneri: an emerging pathogen. Folia Microbiol (Praha). 2020;65: 275–291.

19. Khodavirdipour A, Jamshidi F, Nejad HR, Zandi M, Zarean R. To Study the Anti-cancer Effects of Shigella Flexneri in AspC-1 Pancreatic Cancer Cell Line in Approach to Bax and bcl-2 Genes. J Gastrointest Cancer. 2021;52: 593–599.

20. Schuster HJ, Gompelman M, Ang W, Kooter AJ. An adult case with shigellosis-associated encephalopathy. BMJ Case Rep. 2018;2018. doi:10.1136/bcr-2017-222372

21. Aubin GG, Bémer P, Kambarev S, Patel NB, Lemenand O, Caillon J, et al. Propionibacterium namnetense sp. nov., isolated from a human bone infection. Int J Syst Evol Microbiol. 2016;66: 3393–3399.

22. Corvec S, Guillouzouic A, Aubin GG, Touchais S, Grossi O, Gouin F, et al. Rifampin-Resistant Cutibacterium (formerly Propionibacterium) namnetense Superinfection after Staphylococcus aureus Bone Infection Treatment. J Bone Jt Infect. 2018;3: 255–257.

23. Yasutomi E, Ueda Y, Asaji N, Yamamoto A, Yoshida R, Hatazawa Y, et al. Liver abscess caused by Cutibacterium namnetense after transarterial chemoembolization for hepatocellular carcinoma. Clin J Gastroenterol. 2021;14: 246–250.

24. Corvec S, Fayoux E, Tessier E, Guillouzouic A, Moraru C, Lecomte R, et al. Cutibacterium namnetense osteosynthetic cervical spine infections: experience with two cases. Eur J Clin Microbiol Infect Dis. 2024;43: 395–399.

25. Summanen PH, Durmaz B, Väisänen M-L, Liu C, Molitoris D, Eerola E, et al. Porphyromonas somerae sp. nov., a pathogen isolated from humans and distinct from porphyromonas levii. J Clin Microbiol. 2005;43: 4455–4459.

26. Boutriq S, González-González A, Plaza-Andrades I, Laborda-Illanes A, Sánchez-Alcoholado L, Peralta-Linero J, et al. Gut and Endometrial Microbiome Dysbiosis: A New Emergent Risk Factor for Endometrial Cancer. J Pers Med. 2021;11. doi:10.3390/jpm11070659

27. Xu J, Peng J-J, Yang W, Fu K, Zhang Y. Vaginal microbiomes and ovarian cancer: a

review. American Journal of Cancer Research. 2020;10: 743.

28. Caselli E, Soffritti I, D'Accolti M, Piva I, Greco P, Bonaccorsi G. Atopobium vaginae and Porphyromonas somerae Induce Proinflammatory Cytokines Expression In Endometrial Cells: A Possible Implication For Endometrial Cancer? Cancer Manag Res. 2019;11: 8571–8575.

29. Walther-António MRS, Chen J, Multinu F, Hokenstad A, Distad TJ, Cheek EH, et al. Potential contribution of the uterine microbiome in the development of endometrial cancer. Genome Med. 2016;8: 122.

30. Hokenstad A, Mariani A, Walther-Antonio M. Vaginal detection of Porphyromonas somerae is indicative of endometrial cancer diagnosis. Gynecol Oncol. 2017;145: 76.

31. Butler-Wu SM, Sengupta DJ, Kittichotirat W, Matsen FA 3rd, Bumgarner RE. Genome sequence of a novel species, Propionibacterium humerusii. J Bacteriol. 2011;193: 3678.

32. McDade K, Singla A, Pash D, Bavaro M, De La Houssaye C. Neisseria Cinerea Bacteremia Secondary to a Retropharyngeal Abscess. Cureus. 2021;13: e14217.

33. Barth KR, Isabella VM, Clark VL. Biochemical and genomic analysis of the denitrification pathway within the genus Neisseria. Microbiology (Reading). 2009;155: 4093–4103.

34. Ren J-M, Zhang X-Y, Liu S-Y. Neisseria mucosa - A rare cause of peritoneal dialysis-related peritonitis: A case report. World J Clin Cases. 2023;11: 3311–3316.

35. Tauch A, Fernández-Natal I, Soriano F. A microbiological and clinical review on Corynebacterium kroppenstedtii. Int J Infect Dis. 2016;48: 33–39.

36. Giffen SR, Alby K. The Brief Case: A case of bloodstream infection with Corynebacterium kroppenstedtii in an infant. J Clin Microbiol. 2024;62: e0082023.

37. Wong SCY, Poon RWS, Chen JHK, Tse H, Lo JYC, Ng T-K, et al. Corynebacterium kroppenstedtii is an Emerging Cause of Mastitis Especially in Patients With Psychiatric Illness on Antipsychotic Medication. Open Forum Infect Dis. 2017;4: ofx096.

38. Liu J, Chen X, Zhou X, Yi R, Yang Z, Zhao X. Lactobacillus fermentum ZS09 Mediates Epithelial-Mesenchymal Transition (EMT) by Regulating the Transcriptional Activity of the Wnt/β-Catenin Signalling Pathway to Inhibit Colon Cancer Activity. J Inflamm Res. 2021;14: 7281–7293.

39. Kim M-S, Jo SK, Roh SW, Bae J-W. Alishewanella agri sp. nov., isolated from landfill soil. Int J Syst Evol Microbiol. 2010;60: 2199–2203.

40. Ioannou P, Vougiouklakis G. A Systematic Review of Human Infections by Pseudomonas mendocina. Trop Med Infect Dis. 2020;5. doi:10.3390/tropicalmed5020071

41. Abdelsalam NA, Hegazy SM, Aziz RK. The curious case of Prevotella copri. Gut Microbes. 2023;15: 2249152.

42. Price EP, Sarovich DS, Webb JR, Ginther JL, Mayo M, Cook JM, et al. Accurate and rapid identification of the Burkholderia pseudomallei near-neighbour, Burkholderia ubonensis, using real-time PCR. PLoS One. 2013;8: e71647.

43. Salah ZB, Rout SP, Humphreys PN. Draft Whole-Genome Sequence of the Alkaliphilic Alishewanella aestuarii Strain HH-ZS, Isolated from Historical Lime Kiln Waste-Contaminated Soil. Genome Announc. 2016;4. doi:10.1128/genomeA.01447-16

44. Baris O, Demir T, Gulluce M. Investigation of In vitro Mineral forming bacterial isolates from supragingival calculus. Niger J Clin Pract. 2017;20: 1571–1575.

45. Huse SM, Ye Y, Zhou Y, Fodor AA. A core human microbiome as viewed through 16S rRNA sequence clusters. PLoS One. 2012;7: e34242.

46. Long PA, Sly LI, Pham AV, Davis GHG. Characterization of Morococcus cerebrosus gen. nov., sp. nov. and Comparison with Neisseria mucosa. International Journal of Systematic and Evolutionary Microbiology. 1981;31: 294–301.
